# “Integrative genomics study of microglial transcriptome reveals effect of DLG4 (PSD95) on white matter in preterm infants”

**DOI:** 10.1101/105288

**Authors:** Michelle L Krishnan, Juliette Van Steenwinckel, Anne-Laure Schang, Jun Yan, Johanna Arnadottir, Tifenn Le Charpentier, Zsolt Csaba, Pascal Dournaud, Sara Cipriani, Constance Auvynet, Luigi Titomanlio, Julien Pansiot, Gareth Ball, James P Boardman, Andrew J Walley, Alka Saxena, Ghazala Mirza, Bobbi Fleiss, A David Edwards, Enrico Petretto, Pierre Gressens

## Abstract

Preterm birth places newborn infants in an adverse environment that leads to brain injury linked to neuroinflammation. To characterise this pathology, we present a translational bioinformatics investigation, with integration of human and mouse molecular and neuroimaging datasets to provide a deeper understanding of the role of microglia in preterm white matter damage. We examined preterm neuroinflammation in a mouse model of encephalopathy of prematurity induced by IL1B exposure, carrying out a gene network analysis of the cell-specific transcriptomic response to injury, which we extended to analysis of protein-protein interactions, transcription factors, and human brain gene expression, including translation to preterm infants by means of imaging-genetics approaches in the brain. We identified the endogenous synthesis of DLG4 (PSD95) protein by microglia in mouse and human, modulated by inflammation and development. Systemic genetic variation in *DLG4* was associated with structural features in the preterm infant brain, suggesting that genetic variation in *DLG4* may also impact white matter development and inter-individual susceptibility to injury.

Preterm birth accounts for 11% of all births ^1^, and is the leading global cause of deaths under 5 years of age ^2^. Over 30% of survivors experience motor and/or cognitive problems from birth ^3, 4^, which last into adulthood ^5^. These problems include a 3-8 fold increased risk of symptoms and disorders associated with anxiety, inattention and social and communication problems compared to term-born infants ^6^. Prematurity is associated with a 4-12 fold increase in the prevalence of Autism Spectrum Disorders (ASD) compared to the general population ^7^, as well as a risk ratio of 7.4 for bipolar affective disorder among infants born below 32 weeks of gestation ^8^.

The characteristic brain injury observed in contemporary cohorts of preterm born infants includes changes to the grey and white matter tissues, that specifically include oligodendrocyte maturation arrest, hypomyelination and cortical changes visualised as decreases in fractional anisotropy ^9–13^. Exposure of the fetus and postnatal infant to systemic inflammation is an important contributing factor to brain injury in preterm born infants ^12, 14, 15^, and the persistence of inflammation is associated with poorer neurological outcome ^16^. Sources of systemic inflammation include maternal/fetal infections such as chorioamnionitis (which it is estimated affects a large number of women at a sub-clinical level), with the effect of systemic inflammation in the brain being mediated predominantly by the microglial response ^17^.

Microglia are unique yolk-sac derived resident phagocytes of the brain ^18, 19^, found preferentially within the developing white matter as a matter of normal developmental migration ^12^. Microglial products associated with white matter injury include pro-inflammatory cytokines, such as interleukin-1β (IL1B) and tumour necrosis factor α (TNF-α)^20^, which can lead to a sub-clinical inflammatory situation associated with unfavourable outcomes ^21^. In addition to being key effector cells in brain inflammation, they are critical for normal brain development in processes such as axonal growth and synapse formation ^22, 23^. The role of microglia in neuroinflammation is dynamic and complex, reflected in their mutable phenotypes including both pro-inflammatory and restorative functions ^24^. Despite their important neurobiological role, the time course and nature of the microglial responses in preterm birth are currently largely unknown, and the interplay of inflammatory and developmental processes is also unclear. We, and others, believe that a better understanding of the molecular mechanisms underlying microglial function could harness their beneficial effects and mitigate the brain injury of prematurity and other states of brain inflammation^25, 26^

A clinically relevant experimental mouse model of IL1B-induced systemic inflammation has been developed to study the changes occurring in the preterm human brain ^27, 28^. This model recapitulates the hallmarks of encephalopathy of prematurity including oligodendrocyte maturation delay with consequent dysmyelination, associated magnetic resonance imaging (MRI) phenotypes and behavioural deficits. Here, we take advantage of this model system to characterise the molecular underpinnings of the microglial response to IL1B-driven systemic inflammation and investigate its role in concurrent development.

In preterm infants MRI is used extensively to provide in-vivo correlates of white and grey matter pathology, allowing clinical assessment and prognostication. Diffusion MRI (d-MRI) measures the displacement of water molecules in the brain, and provides insight into the underlying tissue structure. Various d-MRI measures of white matter have been associated with developmental outcome in children born preterm ^29–32^, with up to 60% of inter-individual variability in structural and functional features attributable to genetic factors ^33, 34^. White matter abnormalities are linked to associated grey matter changes at both the imaging and cellular level ^10, 35, 36^, with functional and structural consequences lasting into adulthood ^37, 38^. Tract Based Statistics (TBSS) allows quantitative whole-brain white matter analysis of d-MRI data at the voxel level while avoiding problems due to contamination by signals arising from grey matter ^39^. This permits voxel-wise statistical testing and inferences to be made about group differences or associations with greater statistical power. TBSS has been shown to be an effective tool for studying white matter development and injury in the preterm brain ^40^, providing a macroscopic in vivo quantitative measure of white matter integrity that is associated with cognitive, fine motor, and gross motor outcome ^11, 41, 42^.

In this work we take a translational systems biology approach to investigate the role of microglia in preterm neuroinflammation and brain injury. We integrate microglial cell-type specific data from a mouse model of perinatal neuroinflammatory brain injury with experimental ex vivo and in vitro validation, translation to the human brain across the lifespan including analysis of human microglia, and assessment of the impact of genetic variation on structure of the preterm brain. We add to the understanding of the neurobiology of prematurity by: a) revealing the endogenous expression of DLG4 (PSD95) by microglia in early development, which is modulated by developmental stage and inflammation; and b) finding an association between systemic genetic variability in DLG4 and white matter structure in the preterm neonatal brain.

## RESULTS

### Global effects of systemic IL1B on mouse microglial transcriptome

We first asked how encephalopathy of prematurity affects microglial gene transcription during development, using a previously validated mouse model of IL1B-induced mild systemic inflammation and subsequent neuroinflammation ^27^. In this model IL1B is administered intra-peritoneally from postnatal days 1-5 (Figure 1, Panel A) and we verified that microglia were the predominant myeloid cell in the brain, outnumbering neutrophils, monocytes, macrophages by a factor of 10-100 fold (Supplementary Figure 1, panel a, b). We also observed that the blood-brain barrier remained appreciably intact, with increased expression of tight junction and adherens genes associated with the control of permeability of the blood brain barrier (BBB) (reviewed in ^43^) (Supplementary Figure 2, panel a, b). CD11B+ cells were isolated for analysis by magnetic activated cell sorting (MACS) at postnatal days 1, 5, 10 and 45, and the microglial purity of the cells was verified by FACS and qPCR analysis, demonstrating these cells to be >95% pure microglia (Supplementary Figure 2, panel c-e). These isolated ex-vivo microglia were then used for microarray gene expression analysis.

**Figure 1.**
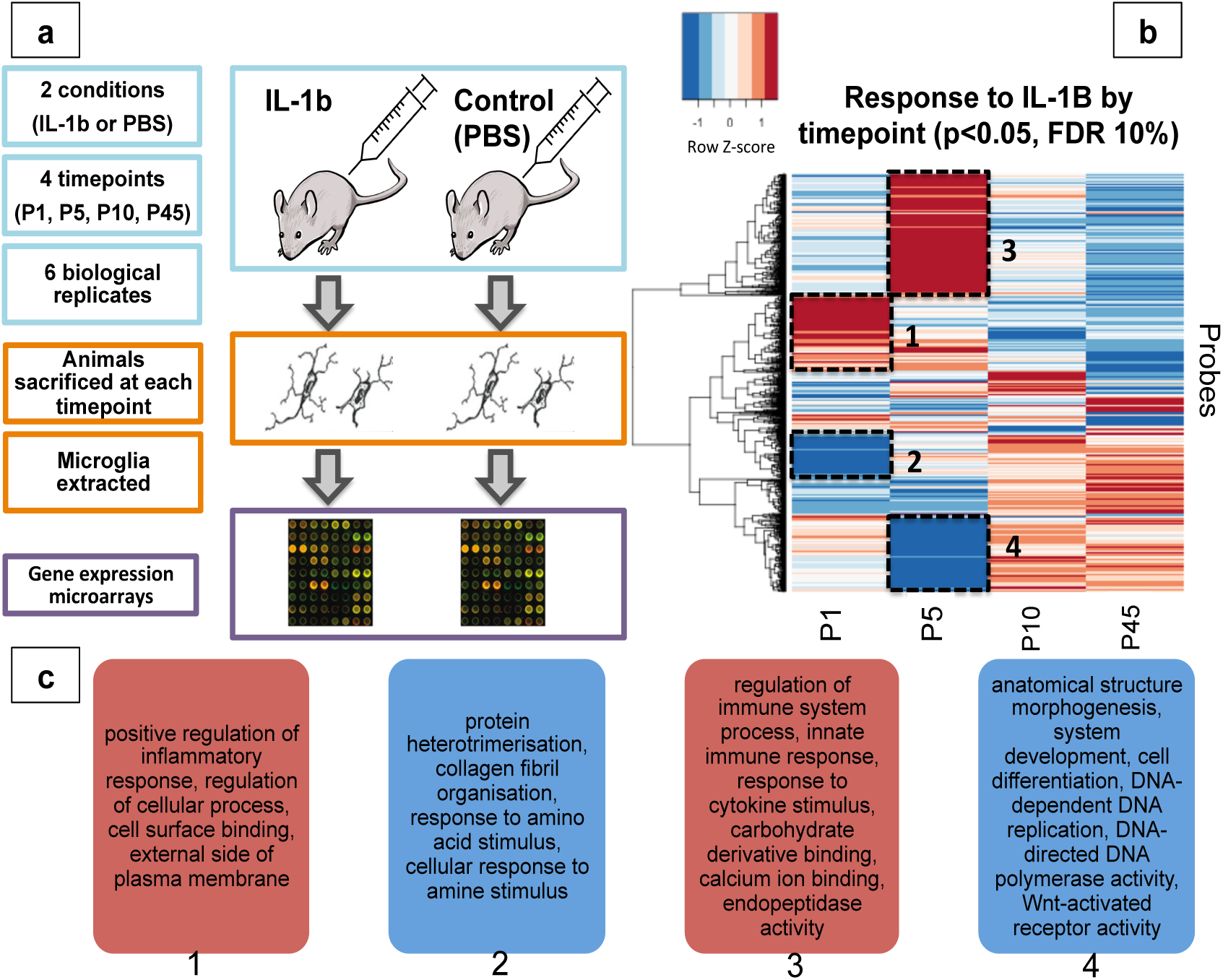
**Overview of in-vivo mouse model, and clustering of expression responses to IL1B by timepoint, with functional annotation summary**. a) IL1B mouse model experimental setup; b) Clustering of expression profiles in response to IL1B identifies up-and down-regulated clusters at each time-point; c) Summary functional annotation of four main clusters identified in b) Red = up-regulated in IL1B versus PBS (control), Blue = down-regulated in IL1B versus PBS.

We examined the global effects of systemic IL1B on the transcriptome of microglia from three angles, by: a) comparing systemic IL1B exposure with control conditions (*response: IL1B*); b) examining transcriptional changes over time (*response: Development*) and; c) assessing whether there is a transcriptional response to IL1B as a function of time (*response: Interaction*). Within the *Development* response is implicit the possible development of microglia themselves ^44^ as well as broader changes at the level of the organism. The *Interaction* response in the MANOVA analysis was used to understand whether the effect of one independent variable (IL1B exposure) on the dependent variables (mRNA levels) was dependent on the value of the other independent variable (time).

After accounting for multiple testing, we found thousands of genes with altered expression in each of the three responses (Supp. Table 1). Functional enrichment analysis of the differentially expressed genes indicated over-representation of different biological processes and pathways across the three responses analysed (Supplementary Figure 3, Supp. Tables 2 and 3). On the whole, the effects of IL1B stimulation and Development on gene expression indicated interdependency such that cytokine-cytokine receptor interaction, transmembrane signalling and cell adhesion molecules were found to be over-represented in all three scenarios. We then used the REVIGO tool ^45^ to summarise these Gene Ontology (GO) enrichments (Methods), which highlighted a set of representative GO biological processes involved in both IL1B stimulation and Development, including immune system processes, cell adhesion, anatomical structure morphogenesis, regulation of multicellular organismal process, cell surface receptor signalling pathway (Supp. Table 4). These findings suggest that IL1B exposure during development has a pervasive effect on the microglial transcriptome, engaging biological processes and pathways relevant to both immune function and growth.

### Patterns of microglia gene expression in response to systemic IL1B exposure

When gene expression profiles were clustered by their response to IL1B at each time-point (Methods), several gene clusters became apparent based on the magnitude of change in IL1B stimulated microglia compared to controls (Figure 1 Panel B, Supplementary Figure 4), with the most prominent clusters observed at P1 and P5 (Z-score above 1 or below −1, difference between IL1B and control p<0.05, False Discovery Rate (FDR) = 10%, two-sample Welch t-statistics, adjusted with Benjamini & Hochberg FDR controlling procedure ^46^). Functional annotation of the genes in these clusters (Figure 1, Supp. Table 5) revealed that inflammatory response processes were up-regulated at P1 and P5 in two distinct waves: an immediate response at P1 (cluster 1) and a subsequent early response at P5 (cluster 3), the latter being a time-point consistent with human preterm gestational age around 32 weeks ^47^. In contrast, biological processes related to anatomical structure development and DNA replication were downregulated at P1 (cluster 2), whereas cell structure and binding-related processes were downregulated at P5 (cluster 4). We also note an overall neutralisation of this transcriptional pattern by P10 with suggestion of a reversal by P45, which was not quantified further but has not been previously observed and may be linked to existing hypotheses regarding a persistent effect of perinatal inflammation ^16^. These observations suggest a rapid and significant transcriptional response of microglia to IL1B exposure, which prioritises inflammatory functions over growth and could lead to disrupted developmental processes.

### Gene co-expression network analysis

We then sought to further examine the relationships between these genes varying in response to IL1B during development. Gene and protein network-based analyses have previously been shown to uncover biological processes involved in disease ^48–50^, with structural (topology) measures being informative of the underlying biology ^51, 52^. The identification of functionally coherent clusters of genes responding to IL1B suggests underlying gene co-regulation, which we sought to explore using gene co-expression network analysis. To this end, we applied Graphical Gaussian Models (GGM) ^53^ to the set of differentially expressed genes identified above, revealing co-expression relationships (i.e. gene co-expression networks) emerging in response to IL1B (*IL1B*), over time (*Development*) or with a differential response to IL1B over time (*Interaction*) (Figure 2). First, we observed a distinct gene membership and function between the three networks, which have only 22 genes in common (Supp. Table 6). We therefore examined the structure of the networks in detail, which showed remarkable topological differences (Supp. Table 7). The Development gene network resembles a small world topology (i.e., most genes are not neighbours of one another), with highest clustering coefficient and degree exponent close to 2. In marked contrast, the IL1B response network is much bigger and more homogenous, and although there are genes with high degree (i.e., hub genes with many other genes connected to them) this does not result in the formation of obvious sub-clusters within the network. The Interaction network has a topological structure intermediate to the other two networks (Figure 2 lower panel and Supplementary Video 1). These differences in co-expression network structure suggest disruption of the small world topology observed under normal conditions, with IL1B leading to a widespread transcriptional response akin to a “genomic storm”, as previously noted in human circulating leukocytes following severe inflammatory stress ^54^.

**Figure2.**
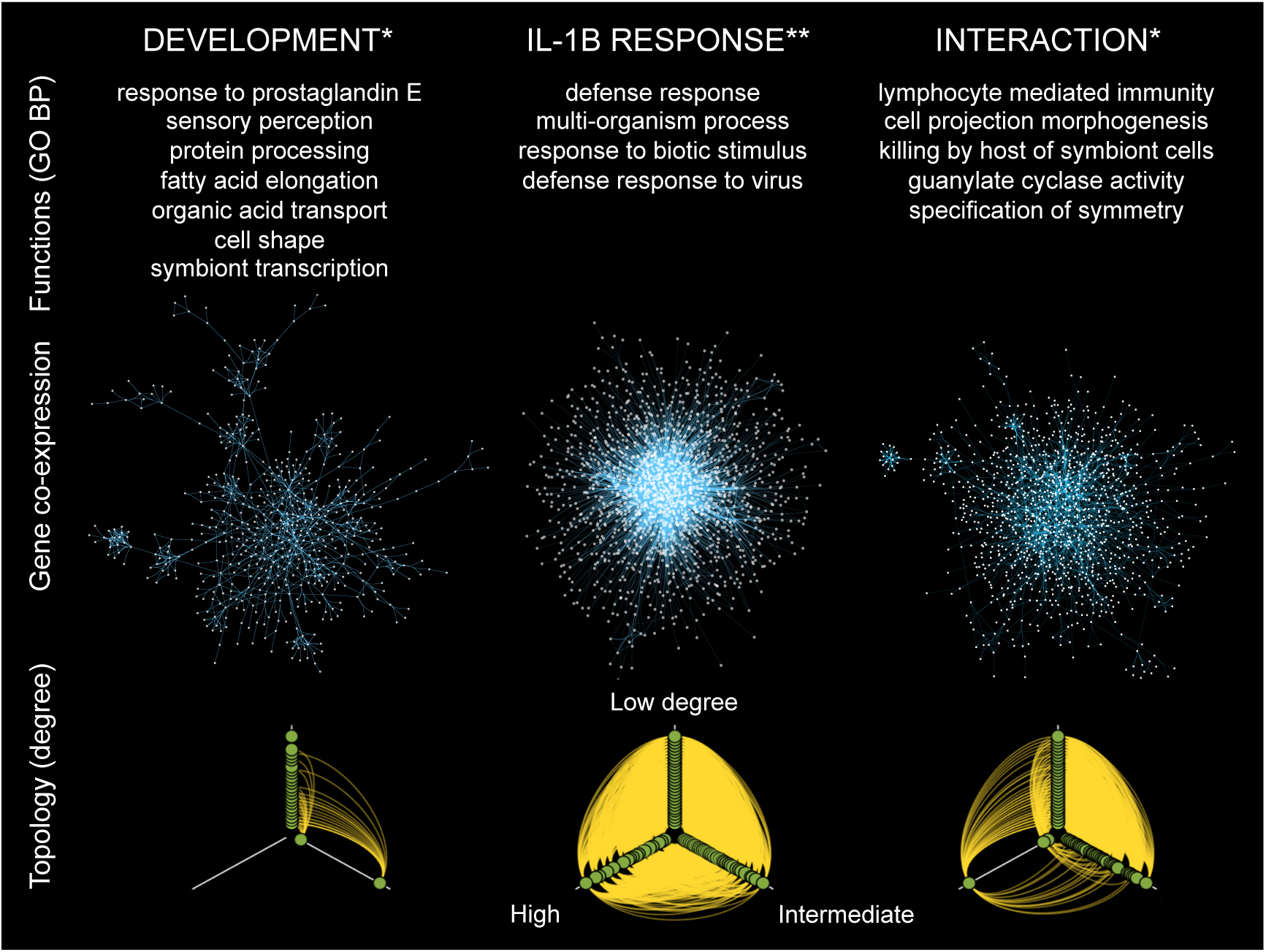
**Gene co-expression networks for three responses: *Development, IL1B and Interaction***. Top: summary of functional annotation. * = nominal p < 0.05, ** = adjusted p < 0.05. Middle: Gene co-expression networks; nodes = genes, edges = partial correlations, local FDR1e^−13^. Bottom: Hiveplots for each network illustrating differences in topology between responses, where nodes are arranged on axes according to their degree; axes ranges clockwise from top: degree < 30, 30≤ degree ≤80, degree x > 80.

Functional enrichment analysis of the three networks suggests that the differences in network topology are paralleled by different biological processes and pathways (Supplementary Figure 5, Supp. Table 8), and there is no overlap between the annotation categories (GO terms or Kyoto Encyclopedia of Genes and Genomes (KEGG) pathways) for the IL1B response and Development networks. The IL1B network is predominantly enriched for GO terms related to defence response, transmembrane signalling and channel activity, whereas both the Development and Interaction networks are point to very broad functional categories. As a result, the network genes are spread among many different categories and the enrichment results mostly did not survive multiple testing correction. Nominally significant functional enrichment terms are therefore included for the annotation of the Development and Interaction networks (Figure 2, top panel). This annotation implies that the structural differences noted earlier between the gene networks are observed also at the functional level, such that the IL1B network has a more specific list of functions linked to immune response, whereas the Development and Interaction networks are involved in broader biological processes.

### Conserved protein-protein interactions and enrichment for neuropsychiatric disease genes

Considering the implications of these co-expression relationships, we asked whether they are conserved at the protein level and whether these are relevant to neuropsychiatric disorders linked to brain development and prematurity. The genes from all three co-expression networks (IL1B, Development and Interaction) were combined into one list and investigated with respect to their shared interactome in the living system. Briefly, we searched for known protein-protein interactions (PPI) within the gene list to identify less redundant network representations and highlight more specific functional interaction processes, an approach which has been previously shown to be useful for disease gene prioritisation ^55^. We used a curated dataset to build a robust PPI network from the genes of interest ^56^ and then carried out a Power Graph Analysis (PGA)^57^ to identify coherent (and simplified) PPI network structures. Power Graphs are lossless representations of graphs based on power nodes (sets of nodes brought together) and power edges (connecting two power nodes, so that all the nodes in the first power node are connected to all the nodes in the second power node). To further simplify and rationalize the Power Graph structures, here we introduce the term super-power node (SPN) to refer to a set of power nodes that form a connected graph. When applied to all genes present in the three co-expression networks, we identified high-confidence connections between 96 proteins. The PGA revealed that 71 of these proteins belong to either one of two main SPNs: SPN1 or SPN2 (Figure 3, Supplementary Figure 6). To independently corroborate the identification of these two SPNs we used a separate approach (DAPPLE)^58^, which interrogates a large database of experimentally derived protein-protein interactions to assess the physical connections among proteins encoded by the genes of interest. In this case, the significance of the PPI derived from the input gene list is assessed empirically (Methods). This analysis replicated the identification of 23/46 edges from SPN1 and 36/41 edges from SPN2, providing further empirical support for the significance of these PPI (permutation-based p-value < 0.001, Supplementary Figure 7). At present, PPIs have rarely been measured in the context of distinct cell types, tissues, or in specific disease conditions, making it challenging to model and understand context-related phenotypes ^59^, but it has also been shown that heterogeneous genomic data contain functional information of protein-DNA, protein-RNA, protein-protein and metabolite-protein interactions ^60–64^. We therefore attempted to gain additional external validation of cell-type specificity for SPN1 and SPN2 by querying the GIANT database of tissue-specific gene networks that includes mapping to tissue and cell-lineage specific functional contexts ^50^. Glia-specific gene interactions within SPN1 and SPN2 were reconstructed with high confidence from prior experimental data (accessed from GIANT database ^50^) (Supplementary Figure 8, panel a), further supporting the consistency and specificity of our findings.

**Figure 3.**
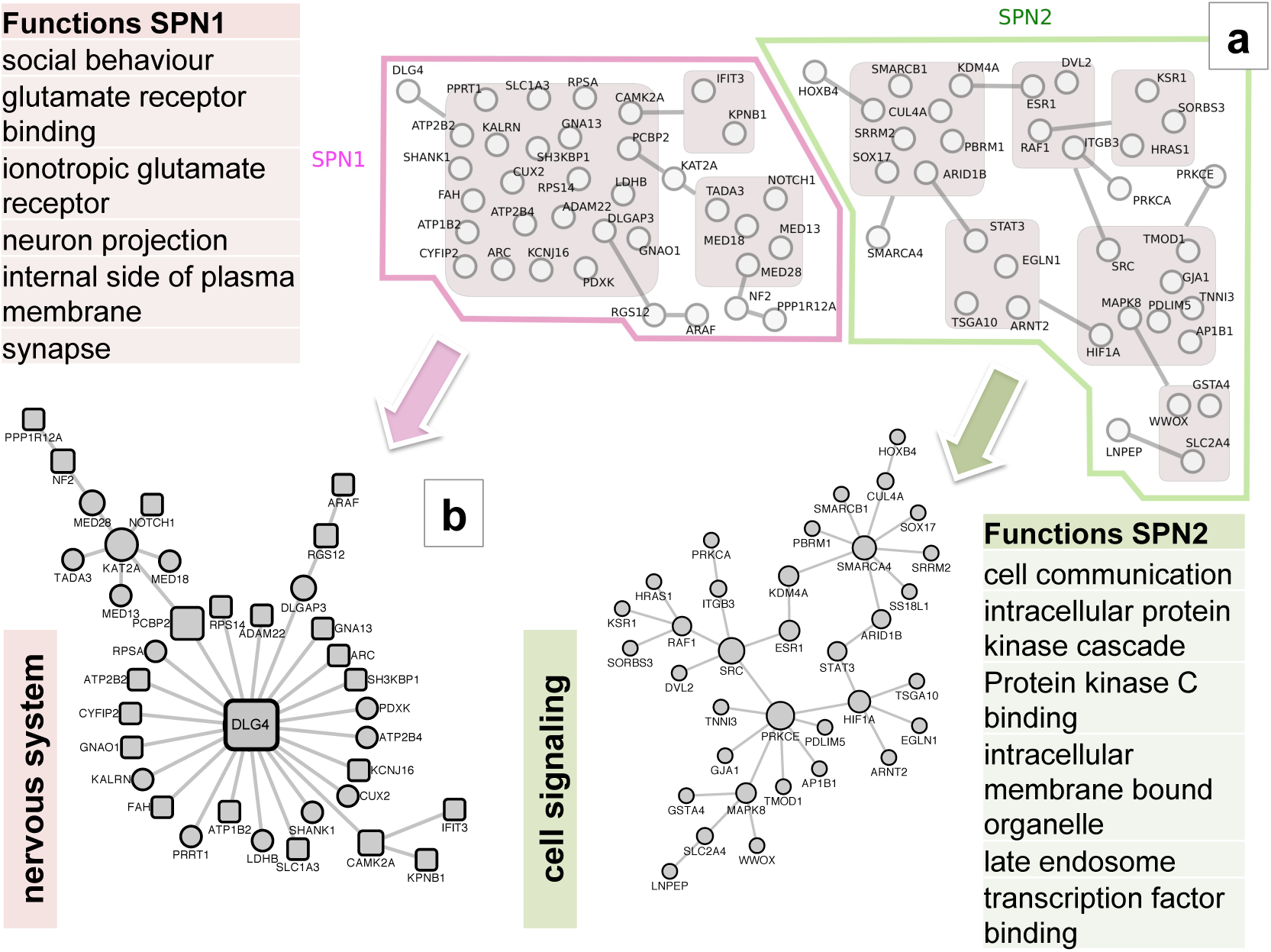
**Protein-protein interactions derived from gene co-expression networks, with grouping into super-power nodes (SPNs) and functional annotation**. a) Two Super Power Nodes (SPNs) in Power Graph Analysis (PGA), showing SPN1 and SPN2 members; nodes = proteins, edges = high confidence curated interactions. Proteins inside grey boxes form modules. Proteins connected to the outline of a grey box are connected to every protein inside that box. b): Conventional visualisation of SPNs. Proteins with square outlines are predicted transcriptional targets of STAT3 transcription factor (TF) at the gene level. Boxes show summary of significant functional enrichment annotation of SPNs (adjusted p-value < 0.01).

Genes coding proteins in the SPNs came from all three gene networks (IL1B response, Interaction and Development) and there was minimal overlap in gene network origin of these proteins (Supplementary Figure 9). This suggests that topologically and functionally different transcriptional networks (Figure 2) might converge to less redundant structures at the protein level (Figure 3), probably reflecting the known modular architecture of PPI networks ^65^ which in our case is reflected in the identification of SPN1 and SPN2. Functional annotation of the two SPNs indicated that these are distinct in function, with SPN1 being significantly enriched for proteins involved in nervous system processes and SPN2 relating to cell signalling and transcriptional regulation (Supp. Table 9, Figure 3). Tissue specificity of expression of SPN genes was queried from the GTEx database ^66, 67^, indicating a pattern of expression of SPN1 genes specific to the brain, whereas SPN2 had a more general tissue distribution (Supp. Tables 10 and 11), in keeping with the protein-based data above.

We then explored whether SPN1 and SPN2 have been involved in brain disorders, and to this aim we tested whether SPN members are enriched for genes annotated for disease association terms using two separate approaches. For this we used the Gene Disease Annotation tool (GDA)^68^, which assesses enrichment of a set of interest (e.g., genes in SPN1) within a set of disease genes (based on Medical Subject Headings categories), with permutation-based significance testing. SPN1 is significantly and specifically enriched for gene-disease links within the Psychology and Psychiatry category, namely Autism and Schizophrenia (p<0.05, 10,000 permutations) (Supp. Table 12). SPN2 has a broader though similar enrichment within the Psychology and Psychiatry category (Supp. Table 12), alongside a very general enrichment across all systems (Supp. Table 13), implying a less specific yet important role in brain disorders. The specific enrichment in SPN1 for genes involved in brain disease was supported by annotation using an independent approach (the WebGestalt tool) ^69^, which prioritises disease-gene links from publications (adjusted p <0.05, Supp. Table 14).

Taken together, these analyses suggest that the gene co-expression relationships observed in microglial cells upon *in vivo* IL1B treatment are at least in part conserved at the protein level; these relationships can be synthetized by two major protein-protein interaction modules (SPNs). Therefore, these SPNs might represent functional modules in protein-protein interaction networks that have been detectable in response to IL1B exposure and development. These SPNs are functionally distinct and enriched for diverse disease gene annotations, with SPN1 specifically enriched for genes involved in neuropsychiatric disorders linked to prematurity, such as *DLG4* in schizophrenia and autism ^70-72^, *SHANK1* in autism ^73^ and *CAMK2A* in several phenotypes ^74^.

### Transcriptional regulation of SPNs

Following the identification of gene co-expression networks and the resulting functional modules at the protein level (SPN1 and SPN2), we set out to examine their potential regulation by transcription factors (TFs) ^75^ and also searched for potential regulatory relationships mediated by TFs involving the members of SPN1 and SPN2. TFs can determine coordinated expression of several target genes (i.e., resulting in a co-expression network) by binding directly to DNA (e.g., at gene promoters), a process that can be mediated by other TFs or proteins that do not themselves interact with DNA directly ^76^.

To identify candidate TFs for the regulation of SPN1-2, we used the PASTAA algorithm ^77^ to analyse the transcription factor binding-site (TFBS) motifs at the promoters of genes that encode for the proteins defining both SPNs (Methods). This analysis across SNP1-2 indicated that a member of SPN2, STAT3 (Signal transducer and activator of transcription 3), as well as other members of the Signal Transducers and Activators of Transcription (STAT) family of TFs (STAT6 and STAT1-alpha), are significantly predicted to bind the promoters of a set of 22/36 (61%) members of SPN1 (p<0.05, Supp. Tables 15 and 16). Given the broad starting point of an unsupervised genome-wide network analysis, we were interested to note that the combined PPI and TF analysis revealed a possible functional relationship between two protein interaction subnetworks. The same relationship was not observed between the STATs TFs and SPN2 genes (Supp. Table 17). The link between the STAT family of TFs and SPN1 genes was also supported by an independent analysis using the Gene Set Enrichment Analysis (GSEA) tool in the Molecular Signatures Database (MSigDB) database (Broad Institute)^78^ to assess the overlap between 615 gene sets (containing genes that share a common TFBS, defined in the TRANSFAC database ^79^) and the genes in SPN1/2. This analysis showed a significant association of members of SPN1 (*Med13, Kcnj16 and Ifit3*) with STAT1 and STAT2 transcription factors (FDR = 2.8%), further supporting a link between SPN2 and SPN1 via the STAT family of TFs.

To seek additional experimental support for the predicted interaction between SPN1 members and the STATs TFs, we investigated the genomic distribution of binding sites of the family of Signal transducers and activators of transcription (STAT1, STAT3, and STAT5) in a publicly available dataset of lipopolysaccharide (LPS)-stimulated primary microglial cultures ^80^. Chromatin immunoprecipitation-promoter microarray data (ChIP-Chip) indicated STAT binding at the promoters of 8/35 (22%) of the genes in SPN1 with either STAT1, STAT3 or STAT5 (Supp. Table 18). On inspection of the STAT3 TF gene expression response to IL1B exposure (Figure 4, panel A), its mRNA level was found to be significantly higher at P1 in microglia exposed to IL1B versus controls, and significantly lower at P5 (FDR = 0.1%). At P1 the mRNA levels of the 22 predicted transcriptional targets of STATs in SPN1 vary, with log_2_ ratios in the range −1.02 to 1.24 (Figure 4, panel b). These results suggest a potential link between a protein interaction network, SPN2, (derived from microglial co-expressed genes in response to IL1B), and a functionally distinct protein interaction network (SPN1) with a putative role in neuronal function and enriched for neuropsychiatric disease genes (Figure 4, panel C). Our transcription factor binding-site (TFBS) analyses indicate that this relationship might be mediated by the STAT3 TF in SPN2 (and potentially by other STAT family members), probably as a result of early *Stat3* gene activation in IL1B exposed cells, as previously reported^81,82^

**Figure 4.**
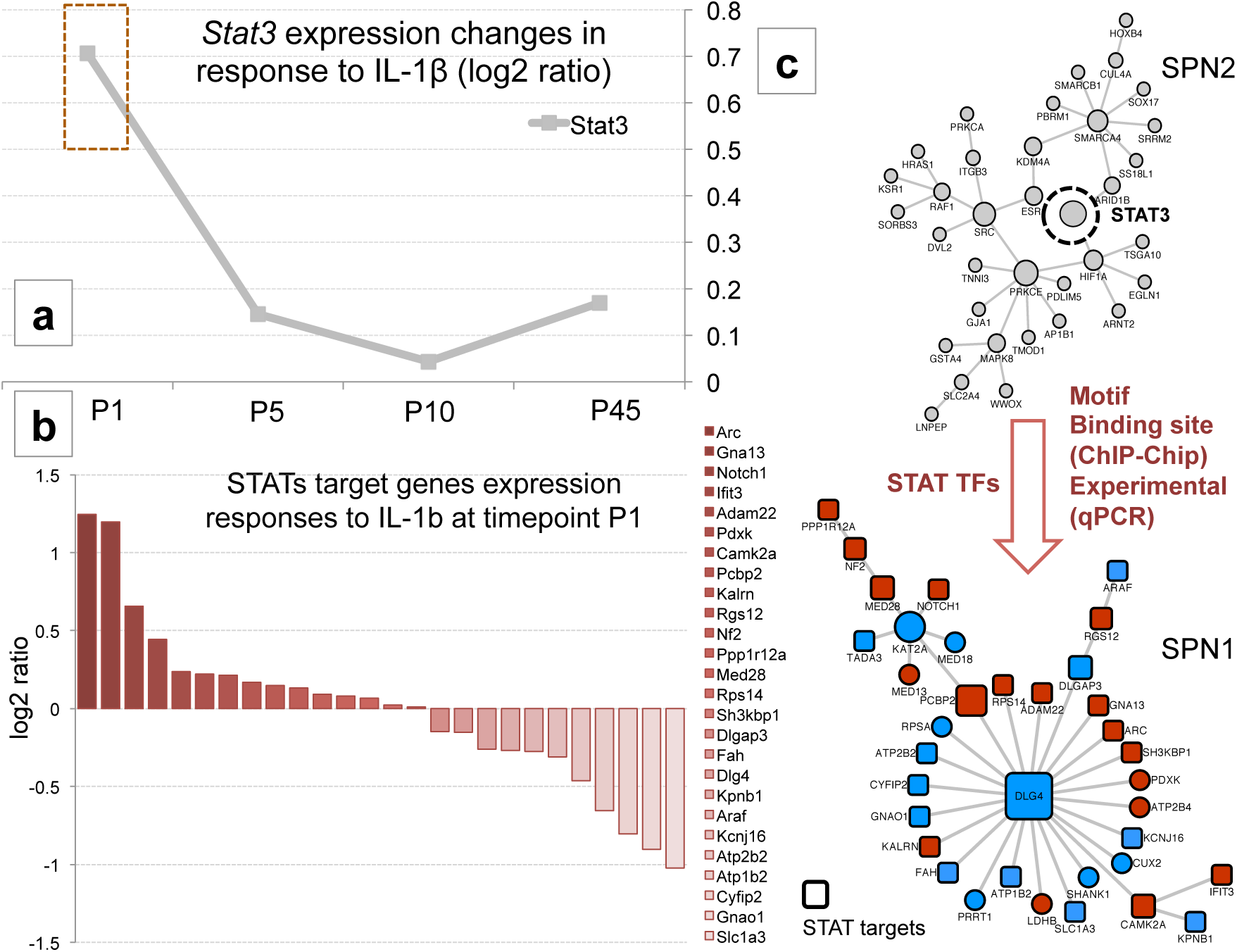
**Transcriptional relationship between STAT3 and SPN1**. a) *Stat3* transcript level by time-point. b) Transcriptional response of target genes of STATs (STAT1, STAT3 or STAT5) to IL1B exposure (log_2_ ratio between IL1B and PBS microarray measures). c) STAT3 from SPN2 is a predicted transcription factor for 22/36 (61%) members of SPN1 (p<0.05, Supp. Tables 15 and 16), corroborated with ChIP-Chip and qPCR data. Red = up-regulated in IL1B versus PBS (control) on microarray, Blue = down-regulated in IL1B versus PBS on microarray.

### Experimental confirmation of effects of STAT3 TF on SPN members

We then tested the validity of our microarray results, and the postulated transcriptional control of SPN1 members by STAT3 TF experimentally. We carried out quantification of gene expression by RT-qPCR on CD11B+ cells MACS isolated at P1 or primary microglia cell cultures from P0-1 mouse cortex and exposed to a vehicle solution or IL1B + IFNg (Methods). A combination of IL1B and IFNg was chosen for two reasons; primarily, that these two cytokines are highly regulated in the brains of IL1B exposed mice (Gressens, unpublished data), making this a useful comparison to the in vivo condition. Second, the combination has been previously demonstrated to cause a moderate but consistent inflammatory reaction in vitro ^83–85^. The transcriptional role of STAT3 in the interaction between SPN1 and SPN2 was investigated by STAT3 pharmacological inhibition followed by RT-qPCR analysis of genes from both SPNs (Methods). MACS-isolated or primary microglia were stimulated by IL1B + IFNg and exposed to vehicle or a small molecule inhibitor of STAT3 (BP-1-102) which binds to the three subpockets of STAT3 SH2 domain and blocks STAT3 phosphorylation, dimerization, and DNA-binding activity ^86^. To investigate the effect of STAT3 inhibition on the SPNs, gene expression was measured by qPCR for a subset of genes, representing a set of general microglia function markers and a subset of genes from SPN1 and SPN2. The profiles and features of the markers that we used were previously characterised by us with in vitro microglia ^83^. Given the high purity of the MACS-isolated CD11B+ cells (>95% microglia, Supplementary Figure 2), we employ them as markers of general microglia activity states and a mechanism of assessing the potential role of the cell rather than for the purposes of cell-type specificity. We used eleven markers broadly grouped as markers of classic pro-inflammatory actions (*Ptgs2, Cd32, Cd86, Nos2)*), immunomodulatory markers (*Il1rn, Il4ra,Socs3, Sphk1*) and regenerative function markers (*Lgal3, Igf1, Cd206*) ^83, 87^, (Methods). IL1B + IFNg induced expression of several pro-inflammatory and immunomodulatory markers and repressed expression of regenerative markers as expected. STAT3 inhibition decreased expression of 3 of the 4 pro-inflammatory markers (*Cd32, Cd86, Nos2),* but had no significant effect on the repression of regenerative markers. By contrast, IL1B + IFNg reduced expression of immunoregulatory marker *Socs3*, while *Sphk1* expression was increased by STAT3 inhibition (Figure 5).

**Figure 5.**
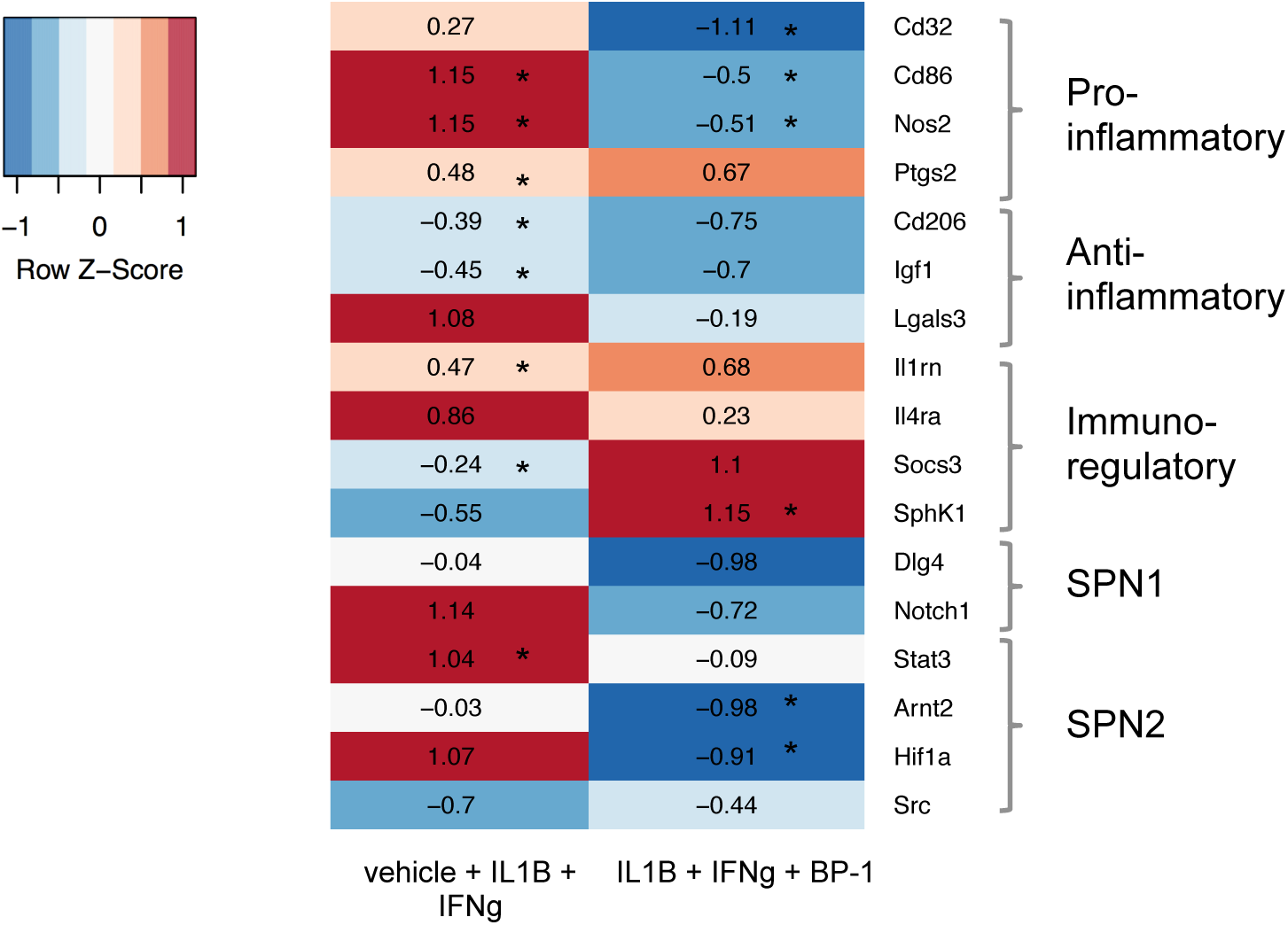
**Effect of STAT3 inhibition (with BP-1) on transcriptional responses to IL1B + IFNγ of SPN genes and microglia functional markers**. MACS-isolated primary microglia (P1 mouse), assessed 4 hours after in-vitro IL1B/IFNy exposure. Heatmap colours represent direction of relative changes (red = increased; blue = decreased). Asterisks in the first column represent significant change in IL1B versus Control, and asterisks in the second column represent significant changes in BP-1 versus IL1B exposure. BP-1: small molecule inhibitor of STAT3 (BP-1-102). vehicle: DMSO (Dimethylsulfoxide, vehicle). IL1B: Interleukin 1 Beta. IFNg: Interferon Gamma. Genes: *Notch1: notch 1; Igf1: insulin-like growth factor 1; Socs3: suppressorof cytokine signaling 3; Stat3: signal transducer and activator of transcription 3; Arnt2: aryl hydrocarbon receptor nuclear translocator 2; Hif1a: hypoxia inducible factor 1, alpha subunit; Nos2: nitric oxide synthase 2, inducible; Cd86: Cd86 antigen; Dlg4: discs, large homolog 4 (Drosophila); Src: Rous sarcoma oncogene; Lgals3: lectin, galactose binding, soluble 3; Ptgs2: prostaglandin-endoperoxide synthase 2; Cd32: Fc fragment of IgG, low affinity IIa, receptor; IL4ra: IL-4 Receptor Subunit Alpha; IL1rn: Interleukin 1 Receptor Antagonist; Cd206: Mannose Receptor, C Type 1; Sphk1: Sphingosine Kinase 1*.

Regarding the analysis of genes from SPN1 (*Dlg4 & Notch1*) we found no effect on *Dlg4* of inflammation or STAT3 inhibition. This lack of response of *Dlg4* gene expression to IL1B is supported by the microarray measurements at P1, which show no significant difference between IL1B and control (PBS) (Figure 4, Student’s t-test p = 0.32). Expression of DLG4 protein in neurons is strongly regulated by post-translational modifications, and in this context our data likely do not provide reliable information on changes at the protein level. *Notch1* gene expression was also not significantly affected by either inflammation or STAT3 inhibition. For the analysis of SPN2 genes, we observed that IL1B + IFNg exposure, significantly increased expression of Stat3, and STAT3 inhibition significantly decreased Arnt2 and HIF1a (Figure 5).

Taken together, these experimental data confirm that IL1B + IFNg exposure induces both pro-inflammatory and immuno-modulatory microglial responses, and that STAT3 presence is required for the pro-inflammatory response and typical modulatory component to occur, as previously suggested in astrocytes ^88^. The transcriptional relationship between SPN1 and SPN2 in the context of inflammation is less clear, leading to the consideration of post-transcriptional mechanisms.

### DLG4 (PSD95): key member of SPN1 and novel role in microglia

We then aimed to clarify how the general IL1B inflammatory stimulus leads to more specific neuronal and neurological effects, and SPN1 was investigated further because of its enrichment for 1) nervous system functional categories and 2) neuropsychiatric disease proteins (reported above). We focused on DLG4 (PSD95), member of the membrane-associated guanylate kinase (MAGUK) family and required for synaptic plasticity associated with neuronal NMDA receptor signalling ^89, 90^, as a key player of interest in SPN1 for further study. This was based on several aspects, including its prominent position as the hub protein of SPN1 (Figure 3), its established neurodevelopmental role in synaptic plasticity^91,92^ and its previously unknown role in microglial function.

These lines of evidence form the hypothesis that the *DLG4* gene plays a role in the neurodevelopmental response to inflammation and may be a potential link to neuropsychiatric disease. These biological processes and subsequent pathological outcome are highly relevant to the preterm population, in which neuroinflammation is posited to contribute to neuropsychiatric sequelae in adulthood ^93^. As such, this allows us to ask whether *DLG4* might be involved in the underlying mechanisms in the human preterm brain.

### DLG4 is present at the mouse microglial membrane and is modulated by development and inflammation

Given our novel finding of endogenous *Dlg4* mRNA expression in mouse microglia by gene expression analysis, we investigated whether the associated DLG4 protein could be observed in mouse microglia, and whether this might be affected by IL1B and development. To investigate the effect of IL1B on the presence of DLG4 in microglia, we assessed immunofluorescence both in tissue sections from pups exposed to IL1B from P1 to P3 and in MACS-isolated CD11B+ cells cultured from P1 mice (Supplementary Figure 11). This allowed us to observe in tissues sections that under control conditions at P1, microglia produce DLG4 protein that is localised to the cell membrane (>95% of IBA1+ cells stained for DLG4) (Figure 6, Postnatal Day 1 and 3 PBS Overlay, Supplementary Video 2). This DLG4 staining disappears by P3, and the protein is still absent from IBA1 positive cells at P45 (data not shown). Following exposure to IL1B however, DLG4 can still be seen at the microglial membrane at P3 along with an apparent change in morphology from a fully ramified to a slightly more amoeboid form (Figure 6, Postnatal Day 1 and 3 IL1B Overlay). However, DLG4 is also not expressed by microglia from IL1B treated mice at P45 suggesting a delay in normative processes as a result of inflammation rather than a permanent change and supporting current hypotheses regarding the neurobiology of prematurity ^94^. These data show an alteration of the normal expression pattern of DLG4 with a possible delay linked to systemic IL1B exposure, and suggest a previously unknown role for DLG4 in the microglial response to inflammation during development.

**Figure 6.**
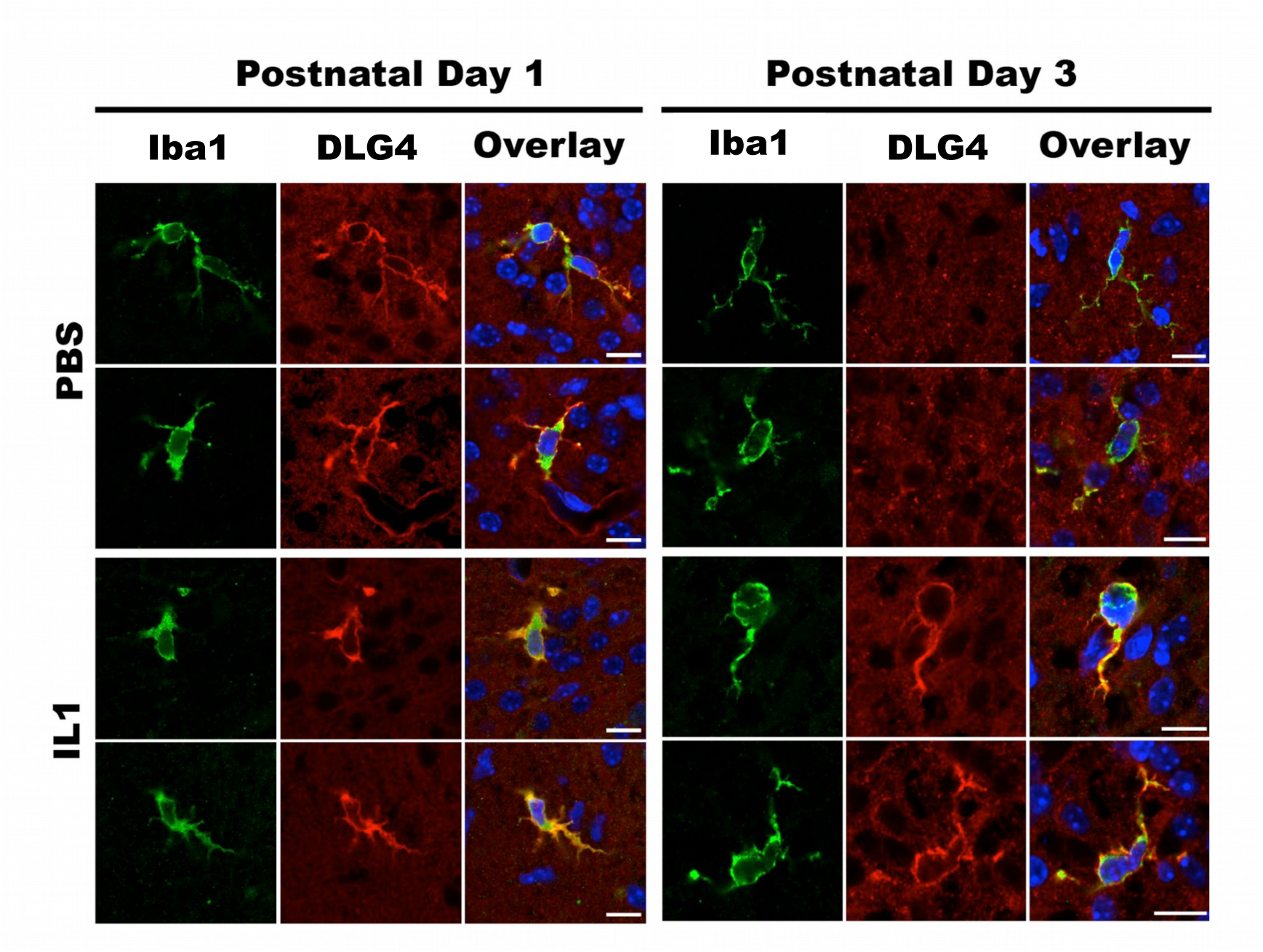
**Immunohistochemistry of mouse brain sections at P1 and P3 with/without in-vivo exposure to IL1B. Double labelling of mouse microglia with DLG4 (PSD95) monoclonal antibodies and a microglial marker (IBA1), under control (PBS) and stimulated (IL1B) conditions, at postnatal day 1 (P1) and P3**.

### Microglia synthesise endogenous DLG4 in early mouse development

To verify that this Dlg4 protein is endogenously synthesised by microglia, rather than phagocytosed neuronal synaptic debris as previously reported later in development ^95^, we stained for DLG4 and Lysosomal Associated Membrane Protein 1 (LAMP1), a marker of lysozymes. No co-localisation of these markers was demonstrable in IBA1+ cells, with LAMP1 immunofluorescence mainly confined to intracellular vesicles and DLG4 consistently and clearly at the microglial cell membrane (Figure 7). A 3D reconstruction of DLG4 and IBA1 immunofluorescence clearly showed membranous staining (Supp. Video 2). In addition, in the developing cortex at P1 and P3 DLG4 protein staining was absent from cells not expressing IBA1. We did observe at P5 staining in a limited number putative cortical neurons DLG4 protein and this timing of neuronal expression is in keeping with the reported normal developmental expression of this protein in neurons in mice and comparative time points in humans ^96^. We also found DLG4 protein at the membrane of MACS-isolated primary mouse microglia maintained ex-vivo for 96 hours, in the absence of neurons (Supplementary Figure 11).

**Figure 7.**
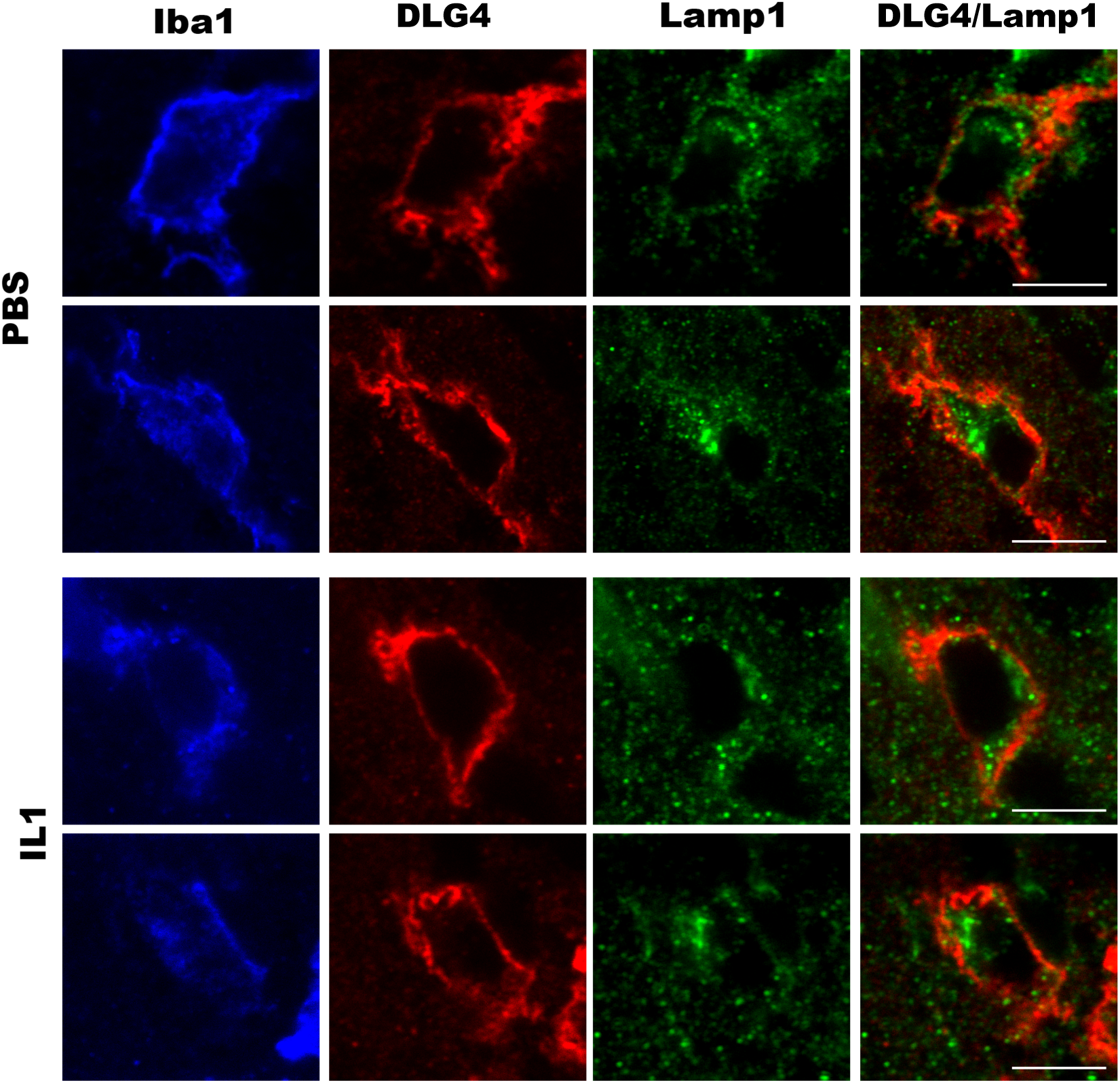
**DLG4 (PSD95) expressed by microglial cells is not localized in lysosomal compartments labelled by LAMP1**. In tissue sections from P1 mouse, microglial cells expressing IBA1 (blue panels) from PBS or IL1B treated animals (P1), DLG4 (PSD95) immunoreactivity (red panels) is predominantly located at the surface of cell bodies and ramifications. Conversely, LAMP1 immunofluorescence (green panels) is mainly confined to intracellular vesicles. No colocalization is thus observed between DLG4 and LAMP1 (DLG4/LAMP-1 panels). Scale bar 5 µm.

### DLG4 is expressed in whole tissues of the human brain

We then asked whether these findings in mice could be translated to human neurobiology, and specifically to microglia in the developing human brain. We first confirmed expression of the *DLG4* gene in the human developing human brain with reference to whole tissue expression data from the Brain Cloud resource ^97^, which allows interrogation of gene expression from human prefrontal cortex in post-mortem samples spanning fetal to adult time-points. We found that the *DLG4* gene is expressed in the human cortex throughout life and that its expression increases rapidly during the first year of life (Supplementary Figure 12, panel A), as previously reported ^98^. We corroborated this observation by accessing *DLG4* cortical gene expression data from the Brainspan Atlas of the Developing Human Brain^99,100^which additionally suggests an apparent decrease in *DLG4* expression at around 21GW, with increasing values from 37GW (term-equivalent age) (Supplementary Figure 12, Panel B). We visualised *DLG4* cortical expression in adults using the Allen Brain Atlas Brain Explorer ^101^ which indicates widespread cortical expression in adulthood (Supplementary Figure 12, Panel C). Previous work has demonstrated that Dlg4 mRNA is targeted for degradation via specific post-translational modifications ^102^, and this may impact its protein distribution during development.

To describe in more detail the regional pattern of expression in the human brain including white matter, we used human brain whole tissue gene expression data from the UK Brain Expression Consortium (UKBEC) database ^103^, which stores paired gene expression and genotype data from 134 brains from normal individuals free of neurodegenerative disorders. We charted *DLG4* gene expression across tissues and found that it is widely expressed in both grey and white matter in adults (Supplementary Figure 8, panel C).

### DLG4 is expressed by microglia in the developing human brain

We then assessed cell-type specific expression of DLG4 protein from microglia of the developing human brain in isolated CD11B cells and in tissue sections. We performed co-localisation of IBA1/DLG4 in MACS-isolated human fetal microglia at 19 gestational weeks (GW) and 21 GW (roughly equivalent to P1 in mouse ^47^) (Methods). We observed that isolated human microglia expressed DLG4 protein together with IBA1 and that this staining appears to increase by activation of microglia to a pro-inflammatory state using lipopolysaccharide (LPS) (data not shown). Further to this, we stained human fetal brain sections through the dorsal cortex at 20, 26 and 30 GW (Methods). We observed a clear co-localisation of IBA1 and DLG4 protein (Figure 8) limited to cells in the proliferative zone and the absence of any other Dlg4 positive cells at 20 GW. At 26 GW we also observed a clear co-localisation of IBA1 and DLG4 protein but also sparse cells in the cortex, putative neurons and at 30 GW we noted very few IBA1 and DLG4 co-localised cells, but many DLG4 positive cells in the cortex with a clear co-localisation of IBA1 and DLG4 protein.

**Figure 8.**
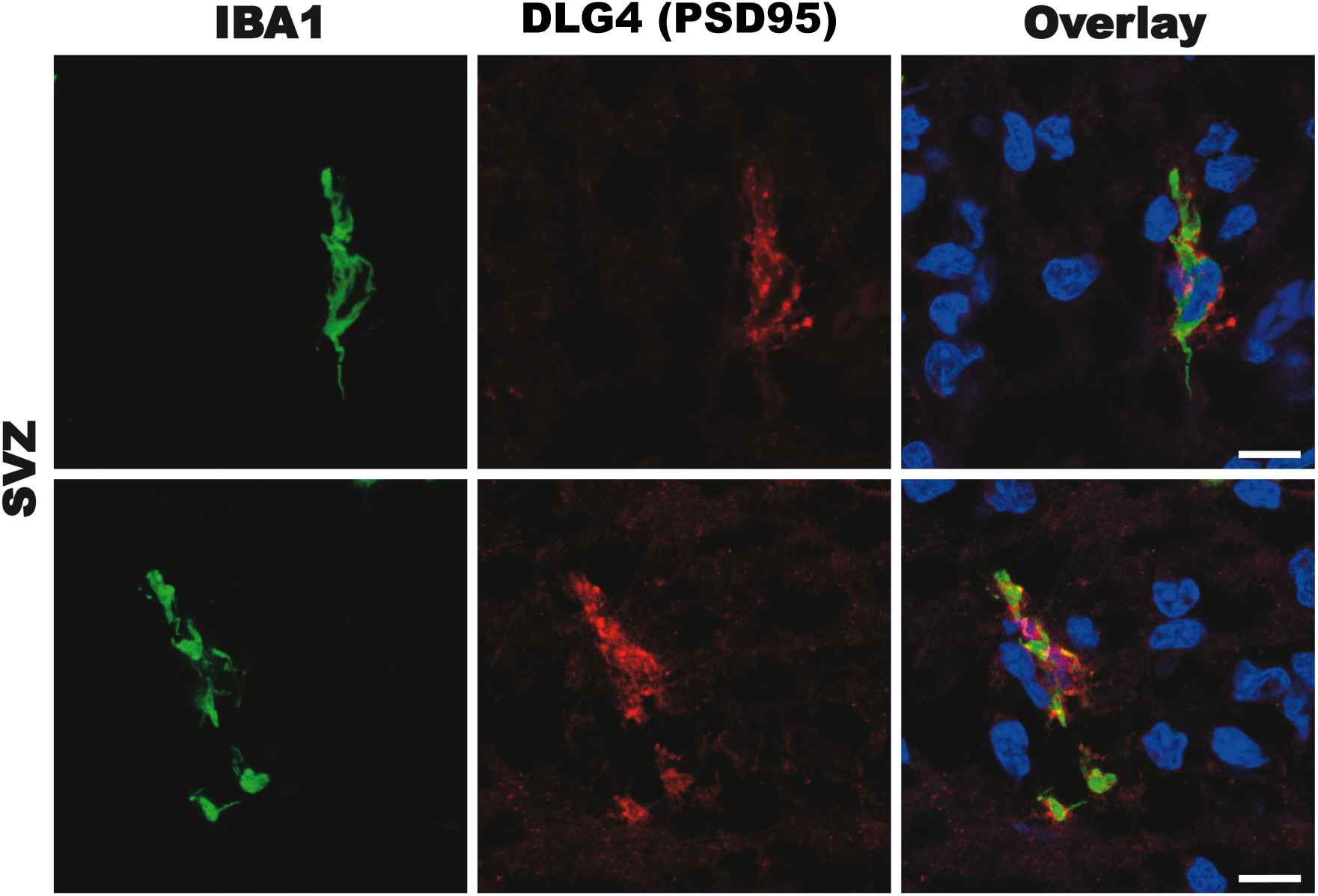
**Immunohistochemistry of human fetal brain sections** Representative photomicrographs from the subventricular zone (SVZ) of sections cut through the dorsal cortex of a 26GW human brain. In green IBA1+ microglia (IBA1) and in red is the DLG4 (PSD95) protein. In the final panel, co-locaisation of IBA1 and DLG4 (PSD95), together with DAPI nuclear staining. Scale bar = 10um.

Taken together, these findings support the widespread expression of the *DLG4* gene in the human brain during development, allied to cell-type specific synthesis of DLG4 protein by human microglia at a time when there appears to be minimal concomitant expression by neurons, which might be further disrupted in the setting of prematurity ^104^.

### Effect of *DLG4* common genetic variants on in-vivo white matter structure in preterm infants

We then set out to examine whether common genetic variants in *DLG4* might have an effect on *invivo* brain features in preterm infants, focusing on white matter given the relevance of this tissue to the preterm neurobehavioural phenotype and its well established diffusion imaging correlation with outcome ^30–32, 105,29^, plus its high content of microglia particularly at this stage of development. MR brain images for two comparable groups of preterm infants (n= 70 and n = 271, Supp. Table 19) were acquired at term-equivalent age, and white matter features (fractional anisotropy, FA) were extracted using Tract-Based Spatial Statistics (TBSS) (Methods). Preterm infant DNA from saliva was genotyped on the Illumina HumanOmniExpress-12 array (Methods), and seven single nucleotide polymorphisms (SNPs) on this array mapped to *DLG4* (dbSNP database ^109^, Supp. Table 20). Of these seven SNPs, only rs17203281 has previously been associated with neuropsychiatric disease. Specifically, the SNP rs17203281 has been found to contribute significantly to prediction of schizophrenia risk in a polygenic non-linear model ^110^, and to form part of a five-marker haplotype associated with schizophrenia ^111^. The same genetic polymorphism has also been associated with changes in cortical regional volume in patients with Williams’ syndrome, which is a well-characterized genetic syndrome overlapping with autism ^71^. This prior information on rs17203281 and our results on *Dlg4* (above) prompted us to investigate the role of rs17203281 in white matter in preterm infants. Infants in both cohorts were categorized by presence or absence of the minor allele (A) for *DLG4* at SNP rs17203281 to test for the effect of minor allele load (MAF in cohort 1 was 0.22 and MAF in cohort 2 it was 0.28), and a general linear model was used to test for a correlation between genotype and white matter FA (Methods). This analysis suggested a significant difference (p < 0.05, FWE-corrected by threshold free cluster enhancement) in FA between infants with or without the minor allele (A) for SNP rs17203281, which was consistent and replicated in both cohorts (Figure 9). There were no significant differences in other clinical features (gestational age at birth, age at scan, days of ventilation) between infants with or without the minor allele (Supp. Table 26). None of the other SNPs at the *DLG4* locus had a replicable effect in both cohorts.

**Figure 9.**
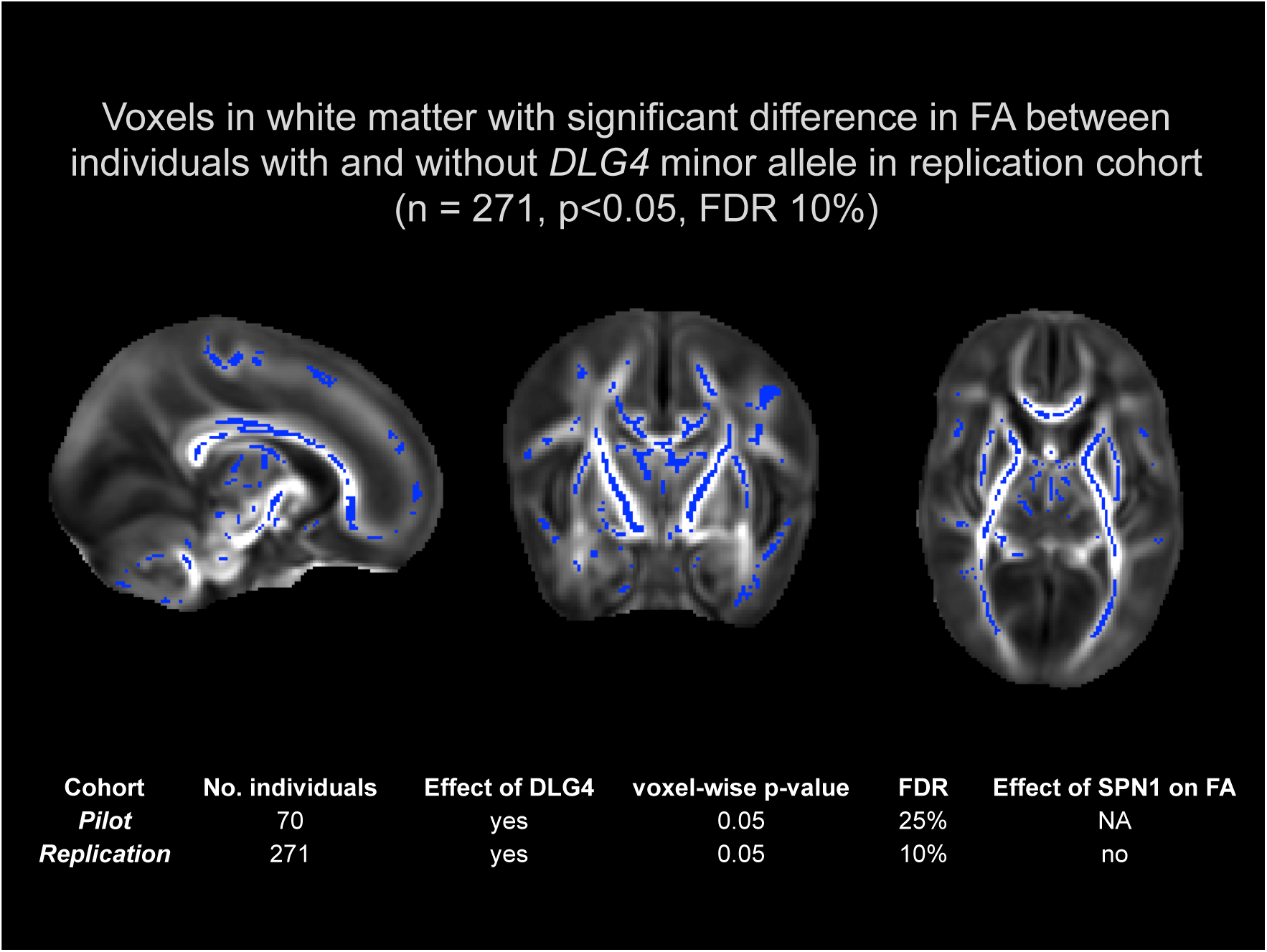
**3T d-MRI brain images for two cohorts of preterm infants acquired at term-equivalent age (cohort 1 = pilot, cohort 2 = replication)**. Images from replication analysis (cohort 2) shown. Views = sagittal, coronal, axial (left-right). Coloured voxels have significantly different diffusion features (fractional anisotropy, FA) between infants with/without the minor allele for *DLG4*. Table inset: Replication of findings in two independent cohorts of preterm infants.

To investigate whether the observed effect of common genetic variation in *DLG4* is specific to this gene or could be extended SPN1 genetic variants as a group, we used a list of all genes in SPN1 as a set of interest and carried out association testing with the phenotype using Joint Association of Genetic Variants (JAG)^112^ in the larger cohort of 271 infants. This tests the null hypothesis of there being no more evidence for association of common genetic variants (SNPs) in SPN1 genes with the phenotype under study than any other random set of an equal effective number of SNPs (Methods). The SPN1-set was tested for association with the real phenotype (query data), and this was repeated with 1,000 permutations of the phenotype (self-contained test) to estimate empirical p-values (see Methods). This analysis showed no significant effect of the SPN1 genes on FA, supporting the specific association of *DLG4* with white matter features in the developing brain rather than a wider and more general effect of the larger SPN1 network.

### eQTL effect of *DLG4* in human brain

We then queried whether the prioritised common genetic variant (rs17203281) in the *DLG4* region might be regulating *DLG4* mRNA expression in the human brain, and so we searched for a possible expression quantitative trait locus (eQTL) effect. To this aim we set out to analyse *DLG4* mRNA expression within individual tissues in the brain, treating the expression levels of the gene as a quantitative trait, so that variations in gene expression that are highly correlated with genetic variation can be identified as expression QTLs (eQTLs). When a genetic variant (SNP) is located within (or is in close proximity) a gene that is significantly associated with the gene’s mRNA variation, this defines a *cis*-acting eQTL ^113^. We returned to the UKBEC database and extracted genotypes for the *DLG4* SNP rs17203281 plus *DLG4* gene expression in the white matter, and carried out an eQTL analysis to link variations in gene expression levels to genotypes (Methods). This analysis of the UKBEC data suggested that normal individuals who are homozygotes for the rs17203281 minor allele (AA) genotype have a significantly higher expression of *DLG4* in white matter than those with the alternate genotypes (AG or GG)(p<0.05), suggesting a *cis*-eQTL effect for rs17203281 (Supplementary Figure 14).

To cross-validate this result we queried the GTEx Portal (www.gtexportal.org/), which provides a searchable resource of multiple different human tissues with genotyping, gene expression profiling, whole genome sequencing and RNA sequencing data ^114^. This revealed that *DLG4* is preferentially expressed in the brain (Supplementary Figure 8, panel B), and has multiple SNPs regulating its expression levels in *cis* (Supp. Table 21), of which one SNP (rs3826408) is in high linkage disequilibrium (i.e., is a proxy SNP)^115^ with rs17203281 (r^2^ = 0.667, D’ = 1). Therefore using separate genetic and gene expression datasets, these analyses suggest *cis*-acting genetic regulation of *DLG4* mRNA expression in the brain, in particular, including the SNP rs17203281 which we previously and independently associated with white matter features in preterm infants.

Overlaps between SPN1/2 genes and disease-associated gene modules in neuropsychiatric disease Rare genetic variants in *DLG4* have previously been associated with autism, schizophrenia and epilepsy in an independent gene network study using exome sequencing data in cohorts of patients with neuropsychiatric disorders ^70^. This identified *DLG4* within a small module of 24 genes involved with synaptic function, in which *de novo* and more severe missense mutations were more likely in individuals with significantly higher intellectual impairment. We investigated the overlaps between the main gene modules reported in ^70^ and our SPN1 and SPN2 genes using a formal test for intersecting gene lists ^116^ (Supp. Table 22, Supplementary Figure 15). We found that there were significant overlaps between SPN1/2 and the reported disease-associated modules. In particular, we found that *DLG4* was the most frequently occurring gene in these overlaps (11/51 occurrences) followed by another SPN1 gene, CAMK2A (4/51). Also of note, our identification of two networks SPN1/2 was broadly captured by the autism Modules 1/2 (reported in ^70^), with *DLG4* is being found in Module 2, considered by the authors to be the synaptic module, in keeping with our interpretation of the broad function of SPN1.

In summary these observations suggest that *DLG4* genetic variation at the level of the organism affects white matter structure in preterm infant, with potential broader implications for associated neuropsychiatric disorders. The latter is suggested by previously published genetic susceptibility data in patients with neuropsychiatric disorders. Our study suggests that the observed differences in white matter features in preterm infants might be mediated by the rs17203281 variant, variation in which is in turn linked to altered gene expression of *DLG4* in the human brain.

## DISCUSSION

In this work, we present a translational systems biology study of the effects of IL1B and development on the microglial transcriptome, in which we integrate mouse and human data from complementary sources to gain a deeper understanding of the neurobiology of preterm brain development and injury. The primary finding of this study is that DLG4 (PSD95) is expressed in microglia during early development and that this developmentally regulated expression pattern is altered by neuroinflammation associated with brain damage in preterm born infants. Specifically, using a mouse model of preterm brain inflammation we analyse cell-type specific patterns of gene co-expression, protein interactions and transcriptional regulation. We identify two networks (SPNs) of interacting proteins, one of which (SPN2) is predicted to interact with the other nervous system-oriented network (SPN1) via the transcriptional regulation by STAT3 of its targets in SPN1. STAT3 TF signalling has previously been linked to microglial immune activity in response to stimuli including lipopolysaccharide (LPS), and the *Stat3* gene has been found to be involved in neuroprotective pathways upregulated in microglia from old mice ^117^. We prioritised DLG4 (PSD95) as a key player given its role as hub of the SPN1 network, its biological importance in neurodevelopment and previously unknown importance in microglia. We document novel dynamics of *Dlg4* mRNA and of DLG4 protein that are endogenous to microglia, and modulated by inflammation and development. In sum, there appears to be a biologically driven discrepancy between *Dlg4* gene expression and DLG4 protein synthesis in response to IL1B, which we hypothesise might be due to post-translational mechanisms modulated by inflammation that have been reported to play an important role in DLG4 protein biology (review in ^102^). We translate these findings to humans by showing evidence of developmentally regulated *DLG4* gene expression throughout human brain tissue, and similarly localise DLG4 protein expression to the membrane of developing human brain microglia. We investigate possible effects of the *DLG4* gene in-vivo in preterm infants, and show a significant impact of common genetic variation in *DLG4* (rs17203281) on preterm human brain imaging features in two independent cohorts. This *DLG4* effect on white matter could be due to measured differences in expression of *DLG4* mediated through a possible *cis*-eQTL action of rs17203281, suggesting that inter-individual genetic variability in *DLG4* gene could affect the response to perinatal inflammation linked to IL1B.

The importance of DLG4 within this biological context could be due to various factors. From a network topology perspective, SPN1 has a star configuration secondary to its central hub DLG4 (where a star consists of a single central node, or network-hub, that is connected to several peripheral nodes). It has been suggested that the star network topology may have specific implications in biological systems, possibly maximising network efficiency at the expense of robustness ^118^ such that perturbing the central node (hub) might have important phenotypic consequences on the whole system ^119^. Hub proteins are important for cellular growth, under tight regulation, continuously evolving ^120^, and more intrinsically disordered than proteins with fewer relations ^121^. Intrinsic disorder refers to regions or whole proteins that do not self-fold intoa fixed 3D structure, resulting in an ensemble of non-cooperatively interchanging structures ^122^. DLG4 is the only member of either SPN1 or SPN2 that qualifies as a hub (under the working definition of a hub as a node with more than 10 connections ^121^), and analysis with the Composition Profiler tool to analyse fractional differences in amino acid composition ^123^ indicated that DLG4 is significantly enriched for disorder promoting amino acids (p < 0.02). From a neurodevelopment point of view, DLG4 is involved in synaptic plasticity ^72, 91, 98^, mediating clustering of both NMDA and potassium channels on the neuronal surface ^89, 90^. At the gene level, a query of the DisGeNET database ^124^, which integrates human gene-disease associations through gene and disease vocabulary mapping, returned consistent association of *DLG4* with neuropsychiatric diseases including schizophrenia and autism (Supp. Table 23) as expected ^125^.

The breadth of the approach used here allows the generation of new hypotheses *in silico* (on the functional role of *DLG4* in microglia), with experimental validation and translation to humans. In this analysis, we integrated unique datasets specific to the biology under study in mice and humans with several open-access data (transcriptomics, protein-protein interactions, transcription factors, genetic susceptibility data and annotations) ^126^ to uncover novel neurobiology of microglia, and linked genetic variation *DLG4* to features of white matter in preterm infants. This study adds to the understanding of the brain injury of prematurity in several ways. At the cellular level we were able to focus on the transcriptional activity of microglia and describe their response to IL1B in great detail, showing two waves of gene expression changes that lead to up-regulation of inflammatory genes and down-regulation of genes involved in growth (Figure 1, Panel C). By using unbiased network analysis approaches we gained a global view of the transcriptome and observed what appears to be a “genomic storm” in response to IL1B ^54^, completely altering the topology seen in development (Figure 2). These dynamics of expression were preserved in interactions at the protein level, giving new insight into the possible roles of well-known genes whose activity has not previously been characterized in this context. In addition, starting from unsupervised network analyses and investigation of relevant protein-protein interaction, we were able to highlight a prominent and yet uncharacterised role for DLG4 (PSD95) in microglia, usually considered an archetypal marker of the neuronal post-synaptic density. Our conclusions on DLG4 from in-depth computational analysis are strengthened by experimental findings: a) Our observation of endogenous cell-type specific expression of *Dlg4* mRNA in mouse microglia; b) DLG4 protein at the microglial cell membrane only in both mouse and human (Figure 6, Supplementary Video 2, Figure 8, Supplementary Figure 11, Supplementary Figure 13). Of specific note, we did not observe DLG4 protein co-localising within vesicles in the cytosol of microglia. Observations of DLG4 protein within microglia as a result of synaptic pruning have been presented at later postnatal ages ^95^, however, before postnatal day 10 we and others ^96^ observe that there is no neuronal expression of DLG4 proteins, due to a well characterised post-translational repression of Dlg4 translation ^96^.

*DLG4* has been previously implicated in neuropsychiatric disease in humans ^70^; our data now extend the repertoire of *DLG4* function in the brain, proposing an association with white matter integrity in the preterm brain, where microglia are known to be predominantly located at this stage (last trimester equivalent)^106, 107, 127^. Given that preterm infants have an increased risk of developing Autism Spectrum Disorders (ASD) and that microglia have been associated with ASD ^128, 129^, we also queried whether behavioural effects of *DLG4* have been investigated in a relevant model. This effect has previously been demonstrated in a mouse knockout model designed to focus on phenotypes relevant to ASD and Williams’ syndrome ^71^. Animals with *Dlg4* deletion (*Dlg4*^-/-^) exhibited increased repetitive behaviour, abnormal communication and social behaviour, impaired motor coordination, and increased stress reactivity and anxiety-related responses, with subtle dysmorphology of amygdala dendritic spines. Investigation of a neural endophenotype of Williams’ syndrome in normal adult humans revealed that rs17203281 homozygote G allele carriers had significantly lesser volume near the right intraparietal sulcus relative to (minor) A allele carriers. Our imaging-genetic analyses suggest a previously unknown contribution of this common genetic variant in *DLG4* to white matter structural features in the preterm infant brain, although we cannot in this study specifically attribute this effect to the action of microglia.

Regarding a potential role for DLG4 protein in microglia and using our in vitro model of microglial activation (exposure to IL1B+ IFNg), we investigated the effects on activation state of blocking the archetypal interaction between the DLG4 protein and the N-methyl-D-aspartate receptor (NMDAR) using a TAT-N-dimer that specifically disrupts the molecular interaction between DLG4 and the NMDAR N2B subunit ^130^. We have previously shown that microglia express functional NMDAR and that these have a moderate but significant effect on microglial activation state and response of the brain to perinatal injury ^131^. We did not see effects of disrupting this relationship on microglial phenotype with any analysis (Methods), namely; RT-qPCR for 12 phenotype markers ^83^, release of chemokines and cytokines (23-plex assay), nitrite nitrate production or phagocytic activity (data not shown). Aside from the archetypal association between DLG4 protein and the NMDAR and its known functions in neurons, we conjecture that the role of DLG4 in microglia in this context may be in the domain of cell-cell communications, such as cross talk between oligodendrocytes, astrocytes, and microglia^132,133^or regulation of glutamatergic and GABAergic signalling, which has been investigated in inflammation and injury ^134–136^. Of interest, DLG4 has been suggested to be involved with the clustering and activity of inwardly rectifying potassium channels (Kir) predominantly expressed in glial cells ^137–139^. Microglial Kir have functional effects on microglial activity and are purported to have possible therapeutic applications in Alzheimer’s disease and Parkinson’s disease ^140, 141^. DLG4 has previously also been observed in oligodendrocytes and proposed to mediate Kir related myelination in development and injury ^138, 142^.

Our observations suggest that under normal conditions microglia make DLG4 protein at P1, which disappears by P3 unless this developmental pattern is disrupted by inflammation, leading to the persistence of DLG4 at the cell membrane. This inflammation-induced persistence has resolved by P45 suggesting a delay rather than permanent change. We suggest that the disappearance of DLG4 protein from the microglial cell membrane between P1-P3 under normal conditions could be to be due to post-transcriptional mechanisms involving the action of micro-RNAs and mRNA degradation^96,143,144^and post-translational mechanisms affecting protein localisation as previously observed in neurons (review in ^102^). Regarding the inflammation-induced persistence of DLG4 protein, we expect this developmental disruption to occur in the human preterm brain, as inflammation is a mediator of the white matter phenotype linked to a delay in oligodendrocyte maturation in these infants ^94^. As such, this forms a novel hypothesis on previously unappreciated biological mechanisms of inflammatory brain injury and supports a deeper future investigation into the exact function of *DLG4* in microglia, and how this might be regulated to reduce perinatal and inflammatory brain injury.

## CONSIDERATIONS AND FURTHER WORK

In this work we use cell-type specific microglia data from a relevant mouse model, as well as cell-type specific data from microglia in developing human brain tissue, plus human white matter genetic variation and gene expression to seek complementary insight into cellular-level neurobiology in perinatal inflammation in infants. These data complement our novel imaging-genomics approach, since genotype data from infants reflect their systemic profiles while current human brain d-MRI analysis methods (including TBSS used here) can only provide insight into structural properties of the brain without cell-type or exact tissue specificity. As such, the relationship observed in the imaging-genomics analysis between *DLG4* variability and brain structure in preterm infants is general, and not attributable specifically to microglia. However, we believe that we have provided a valuable attempt to bridge these conceptual gaps with supporting biological validation.

In focusing our imaging analysis on brain features that best co-localise anatomically with white matter, we hope to capture brain regions that are likely to exhibit the effects of microglial activity since it has been observed that the developing white matter is highly populated with microglia in the perinatal period in both humans and mouse ^106^, and even in adults microglia are typically much more numerous in white matter than in the corresponding regions of neocortex ^108, 145^. It has previously been discussed that TBSS is a reductionist approach to image analysis, and that the specificity and sensitivity of TBSS results are dependent on the quality of the registration and improved by a group-specific target ^146–148^. We have addressed these concerns in detail in the neonatal population by creating an optimised neonatal pipeline ^149^ that includes an initial low degrees-of-freedom linear registration to improve global alignment between neonatal FA maps, and a second registration to a population-average FA map to produce accurate projection of individual data on a skeleton for subsequent multi-subject analysis of white matter diffusivity and anisotropy. In addition we note that our analysis is oriented towards identifying an association between genetic variability and general diffusion properties of white matter rather than attempting a precise spatial localization of this effect within white matter, which would constitute a subsequent specific enquiry making use of higher quality imaging than was available for the current study. We are also currently collecting data to extend a similar approach to the study of grey matter in prematurity.

The model of white matter injury of the preterm infant used here was developed to represent the pathological processes occurring in the contemporaneous human preterm. This model induces a (neuro)inflammatory milieu that captures diverse aspects of neurobiology and behaviour observed in preterm born infants, including; hypomyelination linked to oligodendrocyte maturation arrest, microglial activation, cognitive deficits, decreased fractional anisotropy on MRI, and axonopathy^27^. Nevertheless, as with all animal models, this experimental setup represents an approximation of the human in vivo biology, since an exact emulation of the inflammatory milieu of prematurity is hampered by a lack of specific understanding of these pathological processes in the infant. It is currently under investigation whether this inflammation is triggered by an exposure at a specific time-point or whether there is a persistent inflammatory stimulus and response ^16, 150^. Although this was not a central point of our work and merits investigation in its own right, we attempted to allow for the contribution of longer term effects with the inclusion of the P10 and P45 time-points. Despite these difficulties, the IL1B induced injury model has good clinical comparability and we take supporting data from microglia isolated from this model, cultured human and mouse microglia and human brain tissues. We acknowledge that in this synergy we must also take into consideration limitations that include (but are not limited to) identifying the comparable time points across paradigms and species, and the contribution of other cells to the inflammatory response.

It would also be of interest to delineate formally whether the microglial gene expression response to inflammation also contributes to a robust change in morphology of the cells, and if so whether this structural response can be separated from a specific immune activity or rather if these aspects are inextricably linked. A related point of interest is how much the microglial response to inflammation is conditioned by their own cellular development within the broader context of organismal growth, and how this might differ between the in vivo and in vitro settings in mouse, as well as between mouse and human. The topic of microglial development including how to best characterise it in relation to function remains at present an open question ^44, 106, 151, 152^, and we hope that the availability of these data for future work might help to elucidate further inquiry.

The inflammatory reflex involving the vagus nerve ^153^ may represent an attenuating influence on the IL1B response in vivo, the magnitude and dynamics of which we cannot examine in this work. This is a very interesting balance that appears in fact to converge on modulation of microglia activity via microglial α7 nicotinic acetylcholine receptors (α7AChRs) that limit microglial activation ^154^, possibly signalling via NRG/ErbB4 interactions with DLG4 in microglia ^155–157^ or via endothelial COX2 ^158^.

In terms of possible future therapeutic insights, cell-type specific modulation of dysregulated DLG4 activity in microglia exposed to inflammation could be a strategy to therapeutically attenuate the adverse neurodevelopmental sequelae of prematurity. This approach might be applicable in a precision medicine manner based on inter-individual genetic variation and vulnerability, and we plan to investigate this in future work.

## METHODS

### Animal model

The experimental setup for inducing inflammation-induced white matter injury in the mouse has previously been described in detail ^27^. In brief, a 5µL volume of phosphate-buffered saline (PBS) containing 10µg/kg injection of recombinant mouse IL1B or of PBS alone (control) was injected intra-peritoneally twice a day (morning and evening) on days postnatal (P)1 to P4 and once in the morning on day P5. Animals were sacrificed four hours after the morning injection of IL1B at P1, P5, P10 and P45. For microarray experiments, there were six biological replicates at each time-point which is considered to provide adequate statistical power ^159–161^, and all animals were male Swiss mice (OF1). All *in vivo* and *in vitro* experiments were performed using an alternating treatment allocation. All analyses were performed by an experimenter blinded to the treatment groups.

### Fluorescence-activated cell sorting (FACS) in mouse

Dissociated cells from the cerebrum of mice pups (P1, P3, P5, P10) were centrifuged on a Percoll gradient, as previously described ^162^. Cells were stained using different markers of myeloid cells. Neutrophils were defined as CD11B^hi^LY6G^hi^ whereas monocytes, macrophages and microglia are defined as CD11B^hi^LY6G^lo^. Monocytes were F4/80^lo^ and microglia/macrophages were F4/80^hi^. Finally, microglia were defined as CD45^lo^ and macrophages as CD45^hi^. Thus, cells were stained with anti-CD11B-PerCPCy5.5, LY6G-PE, CD45-FITC, F4/80-APC antibodies (BD Biosciences, NJ, USA). For analysis of purity of CD11B+ microglia MACSing (outlined below), only anti-CD11B-PerCPCy5.5, and F4/80-APC antibodies (BD Biosciences) were used because macrophages in the P1 brain of mice injected with PBS or IL1B represented only 0.6% and 1.7% respectively (Supplementary Figure 1). Cell suspensions were incubated with appropriate dilutions of fluorochrome-conjugated monoclonal antibodies and analysed on a FACS Calibur cytofluorimeter (BD Biosciences). Results were analysed with the Cell Quest Pro software (BD Biosciences). Absolute numbers of different cell populations were calculated by adding 10,000nonfluorescent 10 µm polybead carboxylate microspheres (Polysciences, IL, USA) to each vial, and using the formula: Number of cells = (Number of acquired cells x 10,000)/(Number of acquired beads). Using microspheres and percentages given by the software for each gate, the numbers of different cell populations can thus be obtained.

### CD11B+ microglia magnetic-activated cell sorting (MACS) in mouse

Brains were collected from mice for cell dissociation, and microglia were isolated by magnetic antibody-based cell sorting (MACS) using CD11B antibody according to the manufacturer’s protocol using all recommended reagents and equipment (Miltenyi Biotec, Bergisch Gladbach, Germany) and as previously described ^28^. In brief, mice were intracardially perfused with NaCl 0,9%. After removing the cerebellum and olfactory bulbs, the brains were pooled (per sample at P1, n=3; at P5, n=2; at P10 & P45, n=1) and dissociated using the Neural Tissue Dissociation Kit containing papain enzyme and the gentleMACS Octo Dissociator with Heaters. Brain cells were enriched using the anti-CD11B (microglia) MicroBeads. After elution the isolated cells were centrifuged for 10 minutes at 300g and conserved at -80 °C until RNA extraction or placed into culture and treated as outlined for the primary microglia below. The purity of MACSed CD11B+ fractions was validated using FACS analysis of CD11B fluorescence (described below), and the purity was further validated with qPCR of the positive and negative CD11B cell fractions. (Supplementary Figure 1). Specifically, we used qPCR for glial fibrillary acid protein (Gfap), neuronal nuclear antigen (Neun), Myelin basic protein (Mbp) and Integrin Alpha M (Itgam) gene that encode CD11B. RT-qPCR was performed as described below and analysis confirmed that NeuN, Gfap and Mbp mRNA expression levels were extremely low in CD11B-possitive fraction compare to Itgam mRNA expression. Using CD11B-gated FACS analysis, we observed a slight but significant recruitment of peripheral immune cells - macrophages, monocytes and neutrophils-to the brain over time but no increase in the total number of microglia (Supplementary Figure 1 a, b). However, the relative contribution of these other immune cells to the total pool of CD11B+ cells was 100-1000 fold lower than that of microglia.

### Blood Brain Barrier analysis in rat

The integrity of the blood brain barrier (BBB) was assessed following exposure to either 36 hours (3 injections) or 5 days (9 injections) of twice-daily intraperitoneal injections of IL1B (20µg/kg/injection) in the rat. The phenotype of the injury following IL1B mimics that observed in the mouse and was assessed via gene expression analysis of the total brain and IgG staining of sections of the cerebral cortex as previously described ^28, 163, 164^. Eight animals per group were used for the analysis. For gene expression analysis, pups were sacrificed by decapitation at P2 or P5, 5h after the last injection of IL1B. The brain was removed and the cerebrum was harvested and immediately frozen in liquid nitrogen. Total RNA was isolated using the Rneasy Mini Kit (Qiagen, Courtaboeuf, France) and qPCR performed as previously described ^28^. Genes validated for use as indicators of BBB breakdown were studied ^163^ and primers for each gene measured are listed in Supp Table XD. Data are reported as relative to the reference gene, *Gapdh*. For the analysis of IgG staining, rat pups were killed by decapitation at P5, 5h after the last injection of IL1B. The brains were fixed immediately in 4% formalin and post-fixed for 5 days. Following processing and paraffin embedding, sections were prepared at 10µm and immunolabelling with the antibody anti rat IgG (Sigma, B7139) was performed using the streptavidin-biotin-peroxidase method, as previously described ^164^.

### Microarrays of mouse CD11B+ microglia gene expression and data pre-processing

RNA was extracted and hybridised to Agilent Whole Mouse Genome Oligo Microarrays (8x60K). One biological replicate of IL1B exposure at P1 was removed from the analysis due to low RNA integrity, leaving a total number of 47 samples. Background-corrected log_2_ intensity data were quantile normalised and assigned detection p-values based on level of intensity and proximity to baseline. Genes with an expression p-value <0.05 across all samples were retained for analysis, resulting in a subset of roughly 23 000 genes. Multivariate normality was confirmed with a test for kurtosis implemented in the R package ICS ^165^. Assessment of variance of the data by multi-dimensional scaling (limma package in R^166^) confirmed similar distribution of samples across the IL1B and PBS groups.

### Mouse microglia gene expression response to IL1B and ongoing development

To assess responses gene by gene at each time-point, response to IL1B was assessed by subtracting mean expression values in control PBS from IL1B samples, using hierarchical clustering to group genes by similarity of normalised expression profile, implemented in the heatmap.2 tool in the R gplots package^167^ (Figure 1, Panels B and C, Supp. Table 5, Supplementary Figure 4). Clusters 1-4 were chosen based on magnitude of changes (-1 ≥ Z-≥score1).

Multivariate analysis of variance (maanova R package^168^) was used to evaluate global effects of IL1B exposure and development on gene expression, including an interaction effect of both. Results were filtered to retain changes with p-value < 0.05 (corresponding to a false discovery rate (FDR) 10%^169^) and high coefficient of variation, resulting in three subsets of significantly expressed, highly varying, differentially expressed genes (Supp. Table 1).

### Gene co-expression network reconstruction from mouse microglia

Gene co-expression networks were inferred individually for the Development, IL1B and Interaction responses, using a Graphical Gaussian Models (GGM) implemented within the R software package “GeneNet” ^53, 170^. This computes partial correlations, which are a measure of conditional independence between two genes i.e. the correlation between two genes after the common effects of all other genes are removed. Three separate gene networks were built from the sets of genes identified by MANOVA to show a significant response to IL1B, Development and Interaction effect (Figure 2). Local FDR was set at 1e^-13^ (the most stringent threshold possible, to minimise network size) and a minimum edge-wise partial correlation was set at 0.0075 (the point from which partial correlations increased exponentially across all networks). Hiveplots ^171^ were used to illustrate topological differences between the networks; the axes relate to ranges of node degree and nodes are plotted on each axis according to their degree. An illustrative hiveplot is in Figure 2, with an animation displaying the entire range in Supp. Video 1, and parameters in Supp. Table 25.

### Protein-protein interactions (PPI) and Power Graph Analysis (PGA)

The nodes of all three gene networks (IL1B, Development, and Interaction) were aggregated into one list and used to investigate protein interactions, with both the Netvenn ^56, 172^ and DAPPLE ^58^ tools. The Netvenn tool was also used to perform a Power Graph Analysis (PGA)^57^ on the protein interaction network, and two super-powernodes (SPNs) were identified by examining which sets of powernodes were directly interconnected (Figure 3 and Supplementary Figure 6).

### Functional annotation of gene co-expression and PPI networks

Gene Ontology annotation and enrichment analysis was carried out with the WebGestalt platform ^69^, always using as background the list from which the current set of interest was drawn to avoid inflation of significance e.g. all significantly expressed genes as background for MANOVA genes, and MANOVA genes as background for gene networks.

The REVIGO tool ^45^ was used to summarise GO terms based on semantic similarity measures, using an algorithm akin to hierarchical (agglomerative) clustering. Highly semantically similar GO terms were grouped as guided by the p-values supplied alongside the GO terms. This non-redundant GO term set was visualised using multidimensional scaling to render the subdivisions and the semantic relationships in the data. The disease-gene links used by the Gene Disease Association tool (GDA) ^68^ are assembled from the Genopedia compendium in the HuGE database of Human Genetic Epidemiology ^173^ and the OMIM database ^174^, which are collections of data retrieved from biomedical literature and do not provide cell or tissue specific annotations. While these disease-gene associations are not tissue specific, the relevance and usefulness of protein-protein interaction modules to tissue or cell type specific transcriptional programs has been previously shown by our group and others^64,175,176^

### Transcription factor analysis

The transcriptional control of the protein networks was interrogated in several independent ways. Transcription factor affinities for promoter sequences in the genes coding for the proteins of interest were predicted using the PASTAA tool ^77^. Transcription binding sites for STAT1, STAT3 and STAT5 TFs were identified from experimental ChIP-Chip data of microglia in a P1 rat model with LPS exposure (peaks with FDR <0.2)^80^. Correlation of transcription factor expression levels and potential targets was calculated by finding the linear correlation between each transcription factor and each gene individually.

### Primary in vitro microglial culture in vitro

Primary mixed glial cell cultures were prepared from the cortices of postnatal day (P) 0–1 OF1 mice. Pups of both sexes were included and on average an equal number of males and females were included in each culture. After dissection of the cortices in 0.1 M PBS with 6% glucose and 2% penicillin–streptomycin (PS; Gibco, Cergy Pontoise, France) and removal of the meninges, the cortices were chopped into small pieces and subsequently mechanically dissociated. The suspension was diluted in pre-cooled low glucose Dulbecco’s modified Eagle’s minimum essential medium (DMEM, 31885, Gibco) supplemented with 10% foetal bovine serum (FBS, Gibco) and 0.01% PS. Microglia were isolated from primary mixed glial cultures on day in vitro 14 (DIV14) using a reciprocating shaker (20 min at room temperature) and repeated rinsing with their medium using a 10 mL pipette. Media was subsequently removed, microglia pelleted via centrifugation (300g 10 min) and following re-suspension maintained in macrophage-SFM(serum free medium) (Gibco) at a concentration of 4x10^5^ cells/mL in 6-well culture plates. Culture purity was verified by immunostaining using cell-type specific antibodies against tomato lectin (microglia), glial fibrillary acidic protein (GFAP; astrocytes) and neuronal nuclear antigen (NeuN; neurons) and revealed a >99% purity of microglia.

### Primary microglia ex-vivo, isolated by MACS

Primary microglia were prepared from the brain of P1 mice pups. Brain tissues were dissociated using the Neural Tissue Dissociation Kit containing papain and the gentleMACS Octo Dissociator with Heaters. Microglia were isolated using anti-CD11B (microglia) microbeads (MACS Technology), according to the manufacturer’s protocol (Miltenyi Biotec, Germany) as described above for the preparation of microarray samples. CD11B+ microglia were pelleted via centrifugation and re-suspended in DMEM F12/PS/10% FBS at a concentration of 5x10^6^ cells/mL. Cells were plated in 12-well plates (1ml/well) for RT-qPCR analysis. For immunofluorescence analysis, cells were plated in µ-Slide 8 Well Glass Bottom (Ibidi, Biovalley, France).

### Treatment of microglia with inhibitors for DLG4 and STAT3

Two days after plating, microglia were treated for 6 to 12 hours with vehicle (DMSO, 10µL/ml of culture media), IL1B at 50 ng/mL + IFNg at 20ng/ml, IL1B at 50 ng/mL + IFNg and 20ng/ml + Bp-1-102 (a STAT3 inhibitor; Merck Millipore, Fontenay sous Bois, France) at 15 or 30 µM, or IL1B at 50 ng/mL + IFNg at 20ng/ml + TAT-N-dimer (an inhibitor of the Dlg4 protein NMDA receptor inreraction; Merck Millipore) at 15 or 30nM. Bp-1-102 and TAT-N-dimer were added one hour before the cytokines. At the end of the treatment period, cells were harvested and mRNA extracted for gene expression analysis, and supernatant were collected for nitrites/nitrates or cytokines/chemokines measurement.

### RNA extraction and quantification of gene expression by real-time qPCR

Total RNA from microglial cell cultures was extracted with the RNeasy mini kit according to the manufacturer’s instructions (Qiagen, Courtaboeuf, France). RNA quality and concentration were assessed by spectrophotometry with the Nanodrop™ apparatus (Thermoscientific, Wilmington, DE, USA). Total RNA (1-2µg) was subjected to reverse transcription using the iScript™ cDNA synthesis kit (Bio-Rad, Marnes-la-Coquette, France). qPCR was performed in duplicate for each sample using SYBR Green Supermix (Bio-Rad) for 40 cycles with a 2-step program (5 seconds of denaturation at 96°C and 10 seconds of annealing at 60°C). Amplification specificity was assessed with a melting curve analysis. Primers were designed using Primer3 software, and sequences and their NCBI references are given in Supp. Table 24. The relative expression of genes of interest (GOI) were determined relative to expression of the reference gene, Glyceraldehyde 3-phosphate dehydrogenase (GAPDH). Analyses were performed with the Biorad CFX manager 2.1 software.

### Multiplex cytokine/chemokine assay

Microglia media harvested at the end of the treatment was centrifuged briefly to remove particulates (300xg for 10 minutes). Cytokine and chemokine levels in the microglial media were measured using a Bio-plex 200 with a 96-well magnetic plate assay according to the manufacturer’s instructions (Biorad laboratories, Marnes la Coquette, France). Cytokine and chemokine measured included IL-1α, IL1B, IL-2, IL-6, IL-10, IL-12 (p70), IL-13, G-CSF, GM-CSF, IFNg, TNFá, CXCL1 (KC), CCL2 (MCP-1) and CCL5 (RANTES). All samples were run in duplicate and data was analysed with the Bio-Plex Manager software.

### Nitrites/nitrates assay

Microglia media harvested at the end of the treatment was centrifuged briefly to remove particulates (300xg for 10 minutes). Nitrite/nitrate content was measured using the nitrate/nitrite colorometric assay kit (Cayman Chemical, Ann Arbor, MI, USA) as directed.

### Phagocytosis assay

Phagocytosis of fluorescently labelled *E Coli* particles by microglia was assessed using the pHrodo Red *E. coli* BioParticles Conjugate (Life Technologies) according to the manufacture’s instructions. In brief, 50,000 primary microglial prepared as described above were plated in 48 well plates and after 12 hours of incubation with IL1B at 50 ng/mL + IFN-at 20ng/ml in the presence or absence of TAT-N-dimer at 30 nM, medium was changed to serum free media containing the recommended suspension of bioparticles. Cells were incubated for five hours, before the particle-containing media was removed, washed twice with serum free media, incubated with a solution of trypan blue for 1 min to quench extracellular fluorescence and 1 ml of serum containing media added to each well. The absorbance of each well (including cell free, bead free and media free controls) was read. To adjust for cell density, following reading of the plate the cells were used in an MTT assay.

### RNA extraction and quantification of gene expression by real-time qPCR

Total RNA from primary microglia was extracted with the RNeasy mini kit according to the manufacturer’s instructions (Qiagen, Courtaboeuf, France). RNA quality and concentration were assessed by spectrophotometry with the Nanodrop™ apparatus (Thermoscientific, Wilmington, DE, USA). Total RNA (1-2µg) was subjected to reverse transcription using the iScript™ cDNA synthesis kit (Bio-Rad, Marnes-la-Coquette, France). qPCR was performed in duplicate for each sample using SYBR Green Supermix (Bio-Rad) for 40 cycles with a 2-step program (5 seconds of denaturation at 96°C and 10 seconds of annealing at 60°C). Amplification specificity was assessed with a melting curve analysis. Primers were designed using Primer3 software, and sequences and their NCBI references are given in Supp. Table 24. The relative expression of genes of interest (GOI) were determined relative to expression of the reference gene, Glyceraldehyde 3-phosphate dehydrogenase (GAPDH). Analyses were performed with the Biorad CFX manager 2.1 software.

### Immunohistochemistry of mouse brain sections and isolated cells

Male mice (OF1 strain; Charles River) subjected to the IL1B induced white matter injury outlined above were deeply anesthetized with sodium pentobarbital 3 hours post IL1B or PBS injection at P1, P3 and P10 and perfused transcardially with 4% PFA in 0.1 M phosphate buffer. Brains were post-fixed in the same fixative solution for 2 hours or overnight at 4°C and cryoprotected with 30% sucrose in PBS at 4°C before inclusion in 7% gelatin, 15% sucrose in PBS and freezing in liquid isopentane at -50°C. 12µm thick coronal sections were cut on a cryostat, placed on glass slides and stored at -20°C until immunofluorescent labelling.

Antibodies used were a mouse monoclonal antibody to detect DLG4 ((6G6-1C9) (Product# MA1-045), Thermo Scientific; 1:500^177^, a goat polyclonal antibody to detect IBA1 Ionized calcium binding adaptor molecule 1 (IBA1) (ab5076, Abcam; 1:400 ^178^, and a rabbit polyclonal antibody todetectLysosomal-associatedmembraneprotein1(LAMP-1)(L1418, Sigma;1/200). Secondaryantibodiesusedwerecyanine3-conjugateddonkeyanti-mouse(Jackson ImmunoResearch Laboratories; 1:500), AlexaFluor-488-conjugated donkey anti-goat (Invitrogen; 1:500) and DyLight-405- conjugated donkey anti-rabbit (Jackson ImmunoResearch Laboratories; 1:500).

For sections mounted on glass slides were rehydrated in PBS and pre-incubated in PBS with 0.2% gelatin and 0.25% Triton X-100 (PBS-T-gelatin) for 15 minutes followed by overnight incubation with primary antibodies (anti-PSD95 and anti-IBA1) diluted in PBS-T-gelatin. The sections were rinsed with PBS-T-gelatin and incubated with secondary antibodies diluted in PBS-T-gelatin for 1.5 hours. Depending on the experiments, sections were then rinsed with PBS and incubated with DAPI diluted in PBS (1:1000) for 5 minutes for counter-staining of cell nuclei. All incubations were performed at room temperature, protected from light in a humidified chamber. Finally, the sections were rinsed with PBS, coverslipped with Fluoromount (Southern Biotech) and stored at 4°C until confocal microscopic analysis.

Microglial cells were permeabilized and blocked for 1 h using PBS/0.1% triton/3% horse serum (HS). Primary antibodies: goat anti-IBA1 (Abcam 1:500) and rabbit anti-DLG4 (PSD95) (Abcam, 1:500) or mouse anti-DLG4 (PSD95) (Thermofischer, 1:100), were applied overnight at 4 °C in PBS/1%HS. Fluorescently conjugated secondary antibody to rabbit IgG Cy3 and to goat IgG Alexa 488, were applied for 2 h at 20–25 °C in a humid chamber (1:500, Invitrogen). DAPI (1/500) was applied for 10 minutes (Supplementary Figure 11).

### MACS isolation and inflammatory activation of human CD11B+ microglia

All human post-mortem tissue (cells and tissues) was acquired with ethical approval at The French Agency of Biomedicine (Agence de Biomédicine; approval PFS12-0011). Written informed consent was received prior to donation of fetal tissue. For the collection of human microglia, postmortem tissue without any neuropathological alterations was acquired within 1 hour of scheduled termination (samples from two individuals, 19 and 21 weeks of amenorrhoea). Brain tissue (4g) was mechanically dissociated using 1ml micropipettor in HBSS with Ca2+ and Mg2+, and a single cell suspension was obtained using a 70µM strainer. Isolated microglia were obtained using anti-CD11B microbeads (MACS Technology), according to the manufacturer’s protocol (Miltenyi Biotec, Germany), as described above. CD11B+ microglia were pelleted via centrifugation and re-suspended in DMEM/PS/10% FBS at a concentration of 5x10^6^ cells/mL. Cells were plated in 12-well plates (1ml/well) for RT-qPCR analysis. For immunofluorescence analysis, cells were plated in µ-Slide 8 Well Glass Bottom (Ibidi, Biovalley, France).

Forty-eight hours after plating, human CD11B+ microglia were treated for 4 hours with DMEM (control) or lipopolysaccharide (LPS) 10 ng/mL diluted in DMEM. For qPCR media wereremoved andplates frozen at -80°C. For immunofluorescence experiments, cells were fixed at room temperature with 4% paraformaldehyde for 20 minutes. Lipopolysaccharide (LPS) was used to stimulate human microglia in order to ensure an inflammatory response, since the concentration of IL1B+IFNg used in the mouse experiments might not have been appropriate and we had too few cells available to perform a dose response. We had confidence that the high potency of LPS would lead to a pro-inflammatory response and this was confirmed by the increased expression of TNFa (Supplementary Figure 13).

### Immunohistochemistry of isolated ex-vivo human microglia and brain sections

For the visualisation of IBA1 and DLG4 in human MACS isolated microglia *ex vivo,* cells were treated and stained as per the protsol for mouse sections outlined above. Human brain sections were obtained from post mortem cases from medical abortions at 20 GW, 26 GW and 30 GW for non neurological diagnosis. Tissue was fixed with 4% paraformaldehyde, frozen and sections cut at 12um. Staining for IBA1 (ab5076) and PSD95 (MA1-045) was performed as for mouse tissues above.

### Confocal microscopy

Immunofluorescent stainings of mouse and human tissues and cells were analyzed using a Leica TCS SP8 confocal scanning system (Leica Microsystems) equipped with 488 nm Ar, 561 nm DPSS, and 633 nm HeNe lasers. Eight-bit digital images were collected from a single optical plane using a 63X HC PL APO CS2 oil-immersion Leica objective (numerical aperture 1.40). For each optical section, triple-fluorescence images were acquired in sequential mode to avoid potential contamination by linkage-specific fluorescence emission cross talk. Settings for laser intensity, beam expander, pinhole (1 Airy unit), range property of emission window, electronic zoom, gain and offset of photomultiplicator, field format, and scanning speed were optimized initially and held constant throughout the study so that all sections were digitized under the same conditions.

### Human brain gene expression

#### Braincloud data

Data on *DLG4* expression in the developing brain were accessed from the Braincloud resource, which includes 30,176 probes on 269 samples across the lifespan (fetal through the aged)^97^ and plotted (Supplementary Figure 12, panel A).

#### Allen Brain Atlas data: developing brain transcriptome and adult brain

Developing Transcriptome data as described in the technical white paper were downloaded for *DLG4*^*179*^ and plotted (Supplementary Figure 12, panel B).

Expression of *DLG4* in the adult brain was visualized in the Allen Brain Atlas Brain Explorer software application (http://human.brain-map.org/static/brainexplorer), in which samples from the cerebral cortex are overlaid on an inflated white matter surfaces for each donor's brain, while samples in the subcortical regions of the brain are represented as spheres below the inflated surfaces.

#### UKBEC paired gene expression/genotype data

Human brain gene expression was queried in a collection of 134 brains from individuals free of neurodegenerative disorders^103^. Regional expression of *DLG4* (probeset mean) was extracted for all areas including white matter (Supplementary Figure 8, panel C). Genotypes for rs17203281 in all individuals were also accessed.

### CD11B+ microglia magnetic-activated cell sorting (MACS) in human

Human post-mortem tissue was acquired with ethical approval at The French Agency of Biomedicine (Agence de Biomédicine; approval PFS12-0011). Written informed consent was received prior to donation of fetal tissue. For the collection of human microglia, post-mortem tissue without any neuropathological alterations was acquired within 1 hour of scheduled termination (samples from two individuals, 19 and 21 weeks of amenorrhoea). Brain tissue (4g) was mechanically dissociated using 1ml micropipettor in HBSS with Ca2+ and Mg2+, and a single cell suspension was obtained using a 70µM strainer. Isolated microglia were obtained using anti-CD11B microbeads (MACS Technology), according to the manufacturer’s protocol (Miltenyi Biotec, Germany), as above. CD11B+ microglia were pelleted via centrifugation and re-suspended in DMEM/PS/10% FBS at a concentration of 5x10^6^ cells/mL. Cells were plated in 12-well plates (1ml/well) for qPCR analysis. For immunofluorescence analysis, cells were plated in µ-Slide 8 Well Glass Bottom (Ibidi, Biovalley, France).

### Inflammatory response of primary human microglia ex-vivo

48 hours after plating, microglia were treated for 4 hours with DMEM (control) or lipopolysaccharide (LPS) 10 ng/mL diluted in DMEM. For qPCR media were removed andplates frozen at -80°C. For immunofluorescence experiments, cells were fixed at room temperature with 4% paraformaldehyde for 20 minutes. LPS was used to stimulate human primary microglia in order to ensure an inflammatory response, since the concentration of IL1B+IFNg used in the mouse experiments might not have been appropriate and we had too few cells available to perform a dose response. We had confidence that the high potency of LPS would lead to a pro-inflammatory response and this was confirmed by the increased expression of TNFa.

### Immunohistochemistry of isolated ex-vivo human microglia and brain sections

For the collection of human microglia, post-mortem tissue without any neuropathological alterations was acquired within 1 hour of scheduled termination (samples from two individuals, 19 and 21 gestational weeks (GW), as described above. For the visualisation of IBA1 and DLG4 in human MACS isolated microglia *ex vivo,* slides were permeabilised and blocked for 1 h and primary antibodies, goat anti-IBA1 (ab5076, 1:500) and rabbit anti-PSD95 (MA1-045, 1:500) were applied overnight at 4°C. Fluorescently conjugated secondary antibodies from Invitrogene (1:1000) to rabbit IgG Cy3 and to goat IgG Alexa 488, were applied for 2 h at 20–25 °C in a humid chamber (1:500, Invitrogen).

Samples for sectioning were obtained from a mid-gestation post mortem case under ethical approval at The French Agency of Biomedicine (Agence de Biomédicine; approval PFS12-0011). Specifically, a 26 weeks gestational age case from a spontaneous abortion was identified and the tissue was fixed with 4% paraformaldehyde, frozen and sections cut at 12um. Staining for IBA1 (ab5076) and PSD95 (MA1-045) was performed as for mouse tissues above.

### Imaging data

#### Patient characteristics

*Cohort 1*: Suitable MR images were acquired for 70 preterm infants at term-equivalent age (mean gestational age (GA) 28+4 weeks, mean postmenstrual age (PMA) at scan 40+3 weeks). The cohort consisted of preterm neonates who received care at Queen Charlotte’s and Chelsea Hospital between January 2005 and October 2008, underwent DTI and MR imaging in the neonatal period. Infants were not eligible if they had a chromosomal abnormality, congenital malformation, or congenital infection.

*Cohort 2*: 271 infants (mean GA 29+4 weeks) had suitable imaging at term-equivalent age (mean PMA 42+4 weeks) as part of the EPRIME study (Evaluation of Magnetic Resonance (MR) Imaging to Predict Neurodevelopmental Impairment in Preterm Infants) and were imaged at term equivalent age over a 3 year period (2010-2013) at the Queen Charlotte and Chelsea Hospital, London.

Both studies were approved by the National Research Ethics Service, and all infants were studied following written consent from their parents. All MRI studies were supervised by an experienced paediatrician or nurse. Pulse oximetry, temperature, and heart rate were monitored throughout the period of image acquisition; ear protection in the form of silicone-based putty placed in the external ear (President Putty, Coltene; Whaledent) and Mini-muffs (Natus Medical Inc.) were used for each infant.

#### Image Acquisition

*Cohort 1*: Imaging was performed on a Philips 3-Tesla system (Philips Medical Systems, Netherlands) using an eight-channel phased array head coil. Single-shot echo-planar diffusion tensor imaging (EPI DTI) was acquired in the transverse plane in 15 noncollinear directions using the following parameters: repetition time (TR): 8000 msec; echo time (TE): 49 msec; slice thickness: 2 mm; field of view: 224 mm; matrix: 128 x 128 (voxel size: 1.7531 x 1.753 x 2 mm^3^); b value: 750 sec/mm^2^; SENSE factor: 2. For registration and clinical purposes, a T2-weighted fast-spin echo MRI was also acquired using: TR = 8700 ms, TE = 160 msec, flip angle = 90°, acquisition plane = axial, voxel size = 1.15 x 1.18 x 2 mm, FOV = 220 mm, and acquired matrix = 192 x 186.

*Cohort 2*: MRI was performed on a Philips 3-Tesla system (Philips Medical Systems, Netherlands) using an 8-channel phased array head coil. The 3D-MPRAGE and high-resolution T2-weighted fast spin echo images were obtained before diffusion tensor imaging. Single-shot EPI DTI was acquired in the transverse plane in 32 non-collinear directions using the following parameters: repetition time (TR): 8000 ms; echo time (TE): 49 ms; slice thickness: 2 mm; field of view: 224 mm; matrix: 128 × 128 (voxel size: 1.75 × 1.75 × 2 mm ^3^); *b* value: 750 s/mm^2^. Data were acquired with a SENSE factor of 2. T2-weighted fast-spin echo MRI was also acquired using TR = 8,670 ms, TE = 160 ms, flip angle = 90°, slice thickness = 2 mm, field of view = 220 mm, matrix = 256 × 256 (voxel size = 0.86 × 0.86 × 1 mm^3^).

#### Imaging data selection and quality control

The T2-weighted MRI anatomical scans were reviewed in order to exclude subjects with extensive brain abnormalities, major focal destructive parenchymal lesions, multiple punctate white matter lesions or white matter cysts, since these infants represent a heterogeneous minority (1-3%) with different underlying biology and clinical features to the general preterm population ^180-183^. All MR-images were assessed for the presence of image artifacts (inferior-temporal signal dropout, aliasing, field inhomogeneity, etc.) and severe motion (for head-motion criteria see below). All exclusion criteria were designed so as not to bias the study but preserve the full spectrum of clinical heterogeneity typical of a preterm born population.

Diffusion tensor imaging (DTI) analysis was performed using FMRIB's Diffusion Toolbox (FDT v2.0) as implemented in FMRIB's Software Library (FSL v4.1.5; www.fmrib.ox.ac.uk/fsl) ^184^. Each infant's diffusion weighted images were registered to their non-diffusion weighted (*b*0) image and corrected for differences in spatial distortion due to eddy currents. Non-brain tissue was removed using the brain extraction tool (BET)^185^. FA maps were constructed from 15 or 32 direction DTI, and Tract Based Spatial Statistics ^39^ was used to obtain a group white matter skeleton by using a modified pipeline specifically optimized for neonatal DTI analysis ^149^. Diffusion tensors were calculated voxel wise, using a simple least squares fit of the tensor model to the diffusion data. From this, the tensor eigenvalues and FA maps were calculated, and thresholded at FA > 0.2, then linearly adjusted for PMA at scan and GA.

### Infant genome-wide genotyping

Saliva samples for both infant cohorts were genotyped on the Illumina HumanOmniExpress-12 array, as previously described previously in detail^186^. The genotype matrix was recoded in terms of minor allele counts, including only SNPs with MAF ≥ 5% and ≥ 99% genotyping rate. Seven of these SNPs mapped to the *DLG4* region (Supp. Table 20).

### Association of imaging features with genotype

A general linear model was applied in FSL to test for association between FA values and minor allele count in Cohort 1, and significance was assessed using the randomise tool for nonparametric permutation inference on neuroimaging data^187^ with threshold-free cluster enhancement (TFCE) inference^188^ (Figure 7). This procedure was repeated independently for Cohort 2. There was no significant difference in clinical features (GA, PMA or days of ventilation) between infants with or without the minor allele in either Cohort (Supp. Table 26).

### *DLG4* expression quantitative trait loci (eQTL) analysis

UKBEC genotype and expression data as described above were downloaded for *DLG4* white matter samples and SNP rs17203281. We used the transcript-level expression profile provided in the UKBEC dataset, which is estimated as the Windsorized mean (similar to trimmed mean) of the exon-level probesets that are considered expressed above background noise. It is chosen because it is robust to statistical outliers that may arise from alternative splicing.

Individuals were grouped according to minor allele count 0,1,2 or presence/absence of the minor allele (A) and outliers in the expression value were removed following thresholding by the boxplot.stats function in R. Significant group differences in expression values for the transcription-level were identified using a Student’s t-test (Supplementary Figure 14).

## AUTHOR CONTRIBUTIONS

MLK, EP, PG and ADE designed the research; MLK, JVS, ALS, JY, JA, TLC, ZC, PD, SC, CA, LT, JP, GB, JPB, AJW, BF, PG performed the research; AS and GM contributed genotyping of the replication cohort; MLK, JVS, ALS, JY, JA, TLC, ZC, PD, CA, LT, GB, BF, PG, EP analysed the data; MLK, JVS, JA, BF, ADE, EP, PG wrote the manuscript.

## ACKNOWLEDGEMENTS

Our thanks to the children and families who participated in the study, and the nurses, doctors and scientists who supported the project. This work was supported by grants from Inserm, Université Paris Diderot, Université Sorbonne-Paris-Cité, Investissement d'Avenir (ANR-11-INBS-0011, NeurATRIS), ERA-NET Neuron (Micromet), DHU PROTECT, PremUP, Fondation de France, Fondation pour la Recherche sur le Cerveau, Fondation des Gueules Cassées, Roger de Spoelberch Foundation, Grace de Monaco Foundation, Leducq Foundation, Cerebral Palsy Alliance Research Foundation Australia, Wellcome Trust (WSCR P32674) and The Swedish Research Council (2015-02493). MRI scans of preterm infants were in part obtained in an independent programme of research funded by the National Institute for Health Research (NIHR) Programme Grants for Applied Research Programme (RP-PG-0707-10154.) and in the future a compendium report of the programme will be published in the NIHR Journal. The views and opinions expressed by authors in this publication are those of the authors and do not necessarily reflect those of the NHS, the NIHR, MRIC, CCF, NETSCC, the Programme Grants for Applied Research programme or the Department of Health. In addition, the authors acknowledge financial support from the Department of Health via the NIHR comprehensive Biomedical Research Centre award to Guy's & St Thomas' NHS Foundation Trust in partnership with King's College London and King’s College Hospital NHS Foundation Trust, as well as support from the Medical Research Council (MRC) through Strategic Grant to ADE and Clinical Training Fellowship to MLK (Grant Ref: MR/L001578/1), Duke-NUS Medical School and Singapore Ministry of Health (EP).

